# Primate Astrocyte Evolution Controls the Tempo of Neuronal Development

**DOI:** 10.64898/2026.07.08.736743

**Authors:** Katarzyna Ciuba, Izabela Figiel, Eryk Duński, Garima Virmani, Bondita Dehingia, Misbah Abbas, Keyvan Hemmatvand, Aleksandra Piotrowska, Ewelina Borsuk, Bartłomiej Hofman, Valentina Rava, Elena Taverna, Jakub Włodarczyk, Aleksandra Pękowska

## Abstract

Prolonged neuronal maturation, also referred to as neoteny, constitutes a hallmark of human brain evolution. Yet, the mechanisms controlling neotenic brain development remain poorly understood, and have been defined as neuron-intrinsic. Astrocytes shape synapse formation, activity, and elimination, and have changed substantially between humans and other species. Yet, whether the evolutionary divergence in astrocytes shapes the timing of neuronal maturation is unknown. Here, we show that astrocytes from humans and their closest living relatives, chimpanzees, exert contrasting effects on neuronal maturation: chimpanzee astrocytes accelerate it, whereas human astrocytes delay it, without affecting neuronal survival. Through comparative transcriptomics and epigenomics, we find that evolutionarily reduced APOE expression in human astrocytes underlies the observed delay in neuronal maturation: restoring APOE levels in human astrocytes accelerates neuronal development. We further establish that the Hippo-TEAD signaling represses APOE expression in human astrocytes, revealing a link between the enhanced morphological complexity of human astrocytes and the observed reduced tempo of neuronal development in their presence. Strikingly, neuronal genes differentially impacted by human and chimpanzee astrocytes are associated with schizophrenia, Alzheimer’s disease, and epilepsy, linking astrocyte evolution to disease vulnerability. Altogether, these findings establish that brain neoteny is partly a glial phenomenon, revealing that understanding the pace of human brain development requires understanding how astrocytes, not only neurons, have evolved.

## Introduction

Abstract thinking, language, and creativity constitute the defining features of human cognition, but the cellular and molecular bases of the enhanced mental capacities of our species remain poorly understood. Data gathered over the past years show that neural circuits are expanded in humans compared with non-human primates (NHP)^1–5^. Yet, how this complexity is obtained remains an unsolved fundamental question in evolutionary biology and neurosciences.

Compared to rodents and NHPs, human brain development is markedly prolonged.^6–8^ Post-mortem brain profiling reveales that the periods of synaptogenesis,^9,10^ postsynaptic maturation,^11,12^ synaptic pruning,^13^ and myelination,^14^ are largely supplementary in humans relative to other species^2,6,13,15–18^. Consequently, it has been proposed that evolution endowed our species with enhanced plasticity and the capacity to build a distinctive neuronal architecture^19^ featuring greater complexity by progressively extending the time allocated to brain maturation,.^9,10,13,14^ However, by expanding this critical window of vulnerability, neotenic brain maturation inevitably increased the susceptibility to neurodevelopmental and neuropsychiatric disorders in our species.^13,20,21^ Yet, at present, despite its central role in both cognition and disease, the fundamental question of what determines species-specific differences in neuronal development speed unfolds remains unresolved.

Current evidence shows that neuron-intrinsic, species-specific biochemical reaction rates, protein stability, epigenetic, and metabolic cues constrain the pace of neuronal maturation.^22–33^ This view is further supported by studies showing that human induced neurons mature more slowly than chimpanzee or bonobo neurons even under identical culture conditions, indicating the existence of robust neuron-intrinsic developmental programs.^28,29,34,35^ Critically, all existing models share a fundamental blind spot: they consider neuronal maturation as a cell-autonomous phenomenon, ignoring the fact that neurons mature within a complex cellular environment that actively regulates synapse formation, circuit assembly, and network function. Whether developmental timing is encoded exclusively within neurons or can also be imposed by the cells that surround them remains entirely unknown.

Astrocytes are the principal support and homeostatic glial cells in the brain parenchyma.^36^ They regulate extracellular ion concentration, pH, and supply metabolic substrates and obligatory neurotransmitter precursors to neurons.^36–53^ However, the functions of these unique glial cells extend far beyond homeostasis: astrocytes actively modulate synapse formation, maturation, and elimination, thereby shaping neuronal connectivity and function.^36–53^ Recent studies have further revealed that astrocytes underwent profound evolutionary remodeling, with human cells exhibiting striking increases in size, structural complexity, and transcriptional specialization relative to rodents and other primates.^54–63^ When transplanted into mouse brains, human astrocytes enhance synaptic plasticity and improve learning,^56^ which suggest that changes in astrocyte properties may have contributed to the emergence of human-specific cognitive abilities. Yet, whether astrocyte evolution shapes the pace of neuronal maturation remains unknown.

Here we show that the tempo of neuronal maturation is not determined solely by neuron-intrinsic programs, but is also actively imposed by astrocytes. Using a cross-species co-culture system in which astrocytes from humans, chimpanzees, and rats are paired with rat neurons, we demonstrate that human astrocytes delay neuronal maturation while chimpanzee astrocytes accelerate it. Through comparative transcriptomics and epigenomics, we identify reduced APOE expression as the molecular basis of this delay and uncover elevated Hippo-TEAD signaling as the upstream mechanism that represses APOE in human astrocytes. These findings establish that brain neoteny is partly a glial phenomenon shaped by astrocyte evolution.

### Human and chimpanzee astrocytes impose distinct neuronal maturation states

To test whether evolutionary divergence in astrocytes influences neuronal developmental tempo, we established an interspecies co-culture system in which otherwise identical rat neurons were exposed to astrocytes from different species (**Figure 1a**). We used postnatal day 0 rat hippocampal neurons, which undergo robust excitatory synapse maturation within 14 days in vitro in the presence of rat astrocytes^64,65^ (**Supplementary Figure 1**). Following the elimination of endogenous glia with a mitosis blocker FdU (**Methods**), neuronal maturation became dependent on exogenous astrocyte support (**Supplementary Figure 1**), thereby establishing a defined assay that allowed us to directly compare how astrocytes from different species regulate excitatory synapse maturation and neuronal activity states.

**Figure 1.**
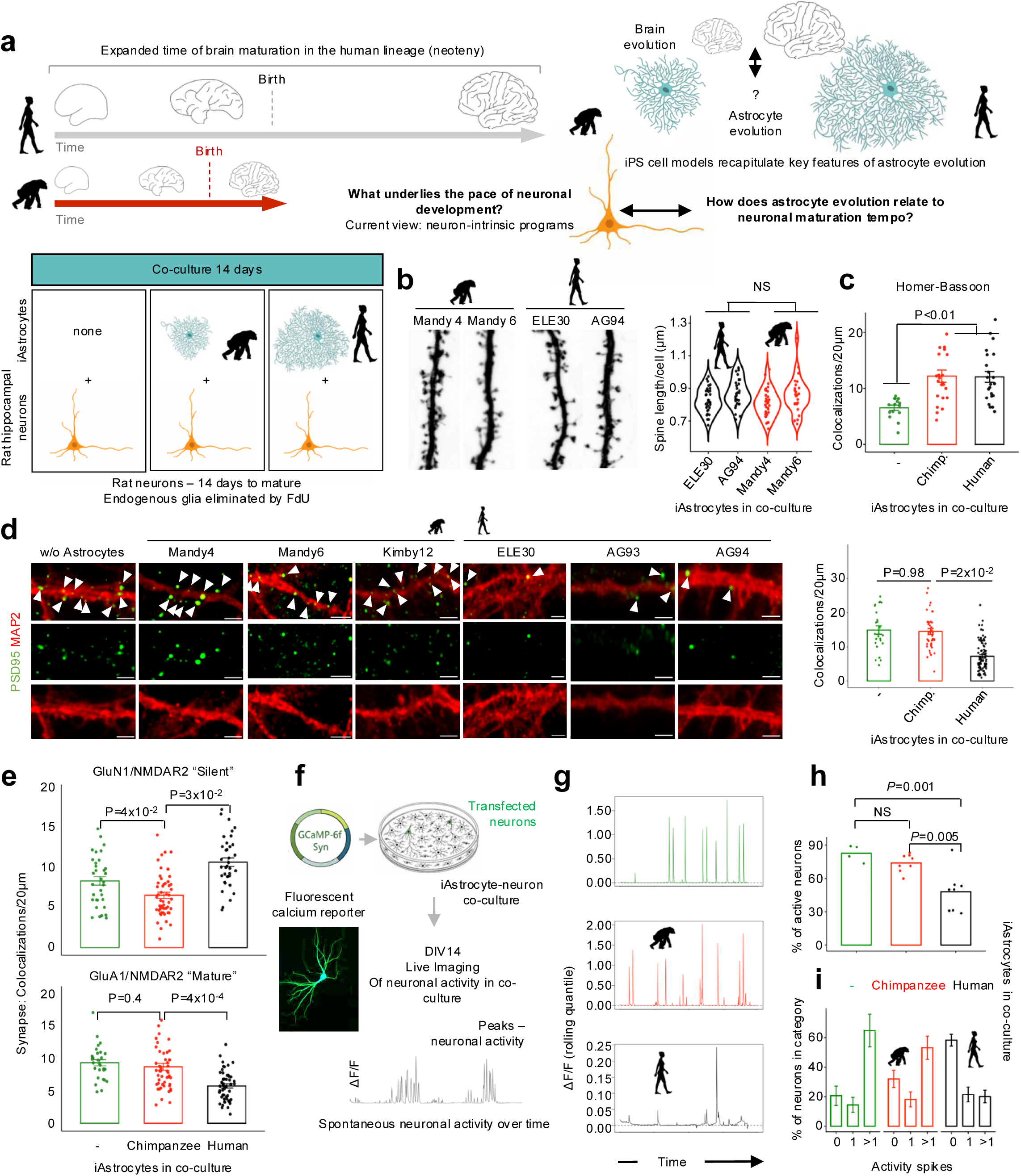
Astrocyte evolution shapes neuronal maturation tempo. **a** Conceptual framework and interspecies astrocyte-neuron co-culture assay. Human brain evolution is characterized by prolonged neuronal maturation (neoteny), which is widely attributed to neuron-intrinsic programs. Human astrocytes underwent substantial evolutionary divergence, including increased morphological complexity and transcriptional specialization, and induced pluripotent stem cell-derived astrocytes (iAstrocytes) recapitulate key features of primate astrocyte evolution. Yet, whether evolutionary changes in astrocytes contribute to the tempo of species-specific neuronal maturation remains unknown. To test this, otherwise identical postnatal day 0 rat hippocampal neurons were co-cultured with human or chimpanzee iAstrocytes, following elimination of endogenous rat glia using 5-fluoro-2′-deoxyuridine (FdU), thereby enabling direct comparison of how astrocytes from different primate species influence neuronal maturation over 14 days in vitro (DIV14). **b** Primate astrocytes do not impact neuronal morphology. Representative images of dendritic segments illustrating spine morphology in GFP-synapsin expressing neurons cultured alone or with chimpanzee or human iAstrocytes (left). Right: quantification of dendritic spine morphology using spine length revealed no significant differences between chimpanzee and human astrocyte co-cultures (*P* > 0.05, Kolmogorov-Smirnov test; n cells = 66 (HS; AG94=35, ELE30=31), 61 (PT; Mandy4=34, Mandy6=27); n spines = 15,232 (HS; AG94=8,171; ELE30=6,461), 12,388 (PT; Mandy4=7,034; Mandy6=5,354).) **c** Human and chimpanzee iAstrocytes promote excitatory synapse formation to a similar extent. Quantification of excitatory synapse density by Homer-Bassoon colocalization. Each dot represents a co-localization within a 20μm fragment of dendrite. N experiments = 3. *P* – value from t-test, after averaging values in each experiment. **d** Excitatory synapses are less mature in rat neurons co-cultured with human iAstrocytes than in the presence of chimpanzee iAstrocytes, as assessed by staining for post-synaptic density 95 marker of maturation and stabilization of excitatory synapses. Representative immunofluorescence images of PSD95-positive mature excitatory synapses (PSD95, green; MAP2, red) in neurons cultured alone or with primate iAstrocytes, with quantification shown on the right. *P* – value from t-test, after averaging values in each experiment. **e** Compared to chimpanzee, human iAstrocytes promote the formation of immature excitatory synapses by rat neurons. Top: quantification of NMDA receptor-containing synapses (GluN1, green; NMDAR2, red) in neurons cultured alone or with either chimpanzee or human iAstrocytes. Bottom: quantification of AMPA receptor-containing synapses (GluA1, green; NMDAR2, red) in neurons co-cultured with chimpanzee or human iAstrocytes. Neurons exposed to chimpanzee iAstrocytes exhibited decreased numbers of immature (GluN1/NMDAR2) and increased numbers of mature (GluA1/NMDAR2) excitatory synapses compared to neurons exposed to human iAstrocytes. Each dot represents a co-localization within a 20μm fragment of a neuronal projection. The indicated statistical differences between species were assessed using a two-sided t-test on the average co-localization inferred in each experiment for each iAstrocyte line (N experiments > 3). Scale bars in all images = 50μm; insets: 2μm. **f** Measurement of neuronal activity in co-culture. Rat neurons were transfected with the genetically encoded calcium indicator GCaMP6f under the synapsin promoter, and spontaneous calcium activity was recorded by live imaging at DIV14. **g** Representative normalized calcium traces from rat neurons cultured alone or in the presence of chimpanzee or human iAstrocytes. Human iAstrocytes suppress spontaneous neuronal activity relative to chimpanzee iAstrocytes. **h** Quantification of the fraction of active neurons in the presence of different primate iAstrocytes. Rat neurons cultured with chimpanzee iAstrocytes exhibited significantly higher spontaneous activity compared to neurons co-cultured with human iAstrocytes. Neurons cultured without astrocytes exhibited activity levels comparable to those in the chimpanzee astrocyte condition. Each dot represents one experiment; at least two cell lines per species were included in each experiment. P-values indicate pairwise comparisons using Student’s t-test. **i** Distribution of neuronal activity states based on the number of spontaneous calcium spikes detected per recording (0, 1, or >1 events). Co-culture with chimpanzee iAstrocytes increased the proportion of neurons exhibiting repeated activity events, whereas human astrocytes increased the fraction of inactive neurons. Data are presented as mean ± SEM; N experiments in d-e = 4

To this end, we generated induced pluripotent stem cell-derived astrocytes (iAstrocytes) from multiple human and chimpanzee lines using our previously established protocol^63^ (**Supplementary Figures 2** and **3**, **Methods**). Astrocytes from both species expressed canonical astrocyte markers, exhibited glutamate uptake, and displayed ATP-evoked calcium responses (**Supplementary Figure 3a-c**). In co-culture, human and chimpanzee iAstrocytes supported neuronal survival to a comparable extent (**Supplementary Figure 3d**), indicating a preserved trophic support across conditions.

We first examined whether primate astrocytes differentially affected neuronal structure and synapse formation. Dendritic spine structure was indistinguishable between conditions (**Figure 1b, Supplementary Figure 4a**), and overall structural synapse number, assessed by Homer-Bassoon colocalization, was similarly preserved across co-cultures (**Figure 1c, Supplementary Figure 4b**). In contrast, neurons co-cultured with chimpanzee iAstrocytes exhibited significantly greater accumulation of post-synaptic density 95 (PSD95), a marker of highly active synapses, than neurons co-cultured with human iAstrocytes (**Figure 1d**; **Supplementary Figure 4c**). This indicates that co-cultures with chimpanzee iAstrocytes promote enhanced postsynaptic maturation compared to those with human iAstrocytes. Next, to directly assess excitatory synapse maturation, we quantified the transition from immature NMDA receptor-only synapses to AMPA receptor-containing mature synapses^66–68^. Neurons co-cultured with human iAstrocytes displayed increased densities of GluN1-positive puncta together with reduced GluA1-positive synapses relative to neurons co-cultured with chimpanzee iAstrocytes (**Figure 1e**; **Supplementary Figure 5ab**), indicating persistence of a more immature excitatory synaptic state of rat neurons exposed to human iAstrocytes. We therefore asked whether these synaptic differences translated into altered neuronal network activity. Using GCaMP6f calcium imaging, which detects spontaneous peaks of calcium concentration reflecting network maturity^69^ (**Figure 1f**), we found that chimpanzee iAstrocytes fostered neuronal activity, whereas neurons co-cultured with human iAstrocytes exhibited markedly reduced spontaneous calcium spikes (**Figure 1g-i**). Human iAstrocyte co-cultures showed both a reduced fraction of active neurons and fewer calcium transients per neuron relative to chimpanzee co-cultures (**Figure 1h-i**; *P* < 5x10^−3^, *ANOVA*).

Together, these findings demonstrate that human and chimpanzee iAstrocytes impose distinct neuronal maturation states. Chimpanzee iAstrocytes promote synaptic maturation and network activity, whereas human iAstrocytes maintain neurons in a less mature functional state. Notably, these effects occur without affecting neuronal morphology, viability or overall synapse formation, indicating that evolutionary changes in astrocyte function selectively modulate the timing of synaptic maturation.

### Human and Chimpanzee Astrocytes Differentially Regulate Neuronal Maturation Programs

To determine how astrocyte species shape neuronal maturation at the transcriptional level, we performed bulk RNA sequencing on co-cultures (**Figure 2a**, ≥3 iPS cell lines per species and ≥3 independent iAstrocyte and neuronal preparations). The sequencing reads were computationally partitioned based on alignment to rat or primate genome (**Methods**),^70^ enabling parallel transcriptional profiling of neurons and astrocytes within the same culture without cell separation.

**Figure 2.**
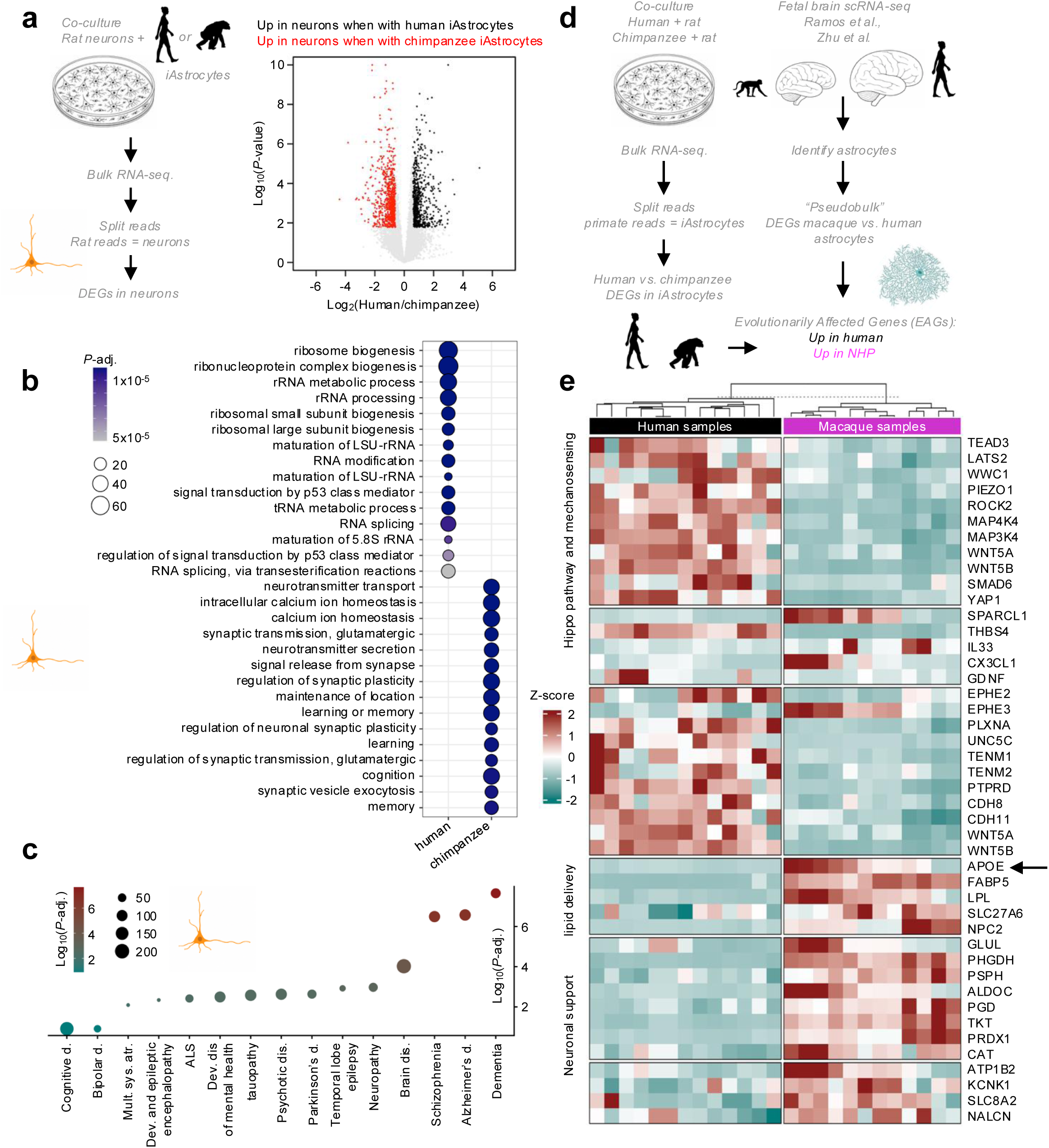
Comparative transcriptomics reveals global changes in neuronal transcriptome in co-culture experiments and identifies evolutionarily diverged astrocyte programs linked to neuronal maturation. **a** The experimental design for transcriptional profiling of neurons in co-cultures. Rat hippocampal neurons were co-cultured with human or chimpanzee iAstrocytes and subjected to bulk RNA sequencing. Sequencing reads were computationally separated to isolate rat- and primate-derived transcripts, enabling differential gene expression analysis specifically in neurons and astrocytes in each co-culture experiment. Differential expression was assessed using *DESeq2* (*design = ∼ batch + species*). Genes with LFC < −0.59 and adjusted *P* < 0.1 (DEG) were used for downstream analysis. **b** Gene ontology enrichment analysis of neuronal genes differentially induced by each astrocyte type. DEGs expressed at higher levels in neurons co-cultured with chimpanzee iAstrocytes were enriched for biological processes related to synaptic transmission, neuronal signaling, learning, and synaptic plasticity. In contrast, DEGs expressed at higher levels in neurons exposed to human iAstrocytes were enriched for processes related to RNA metabolism, ribosomal biogenesis, and transcriptional regulation. **c** Enrichment analysis of neuronal genes differentially induced by chimpanzee astrocytes across gene sets associated with neurological and neuropsychiatric disorders. **d** Strategy for identifying evolutionarily affected genes (EAGs) in astrocytes. Two complementary transcriptomic comparisons were integrated. First, differential gene expression analysis was performed on astrocytes from human/chimpanzee neuron-iAstrocyte co-cultures using bulk RNA-seq. Reads were computationally partitioned by species to separately profile primate astrocytes and rat neurons, and differential expression between human and chimpanzee astrocytes was assessed using *DESeq2* (design = ∼ batch + species). Second, astrocyte transcriptomes from human and macaque fetal brains were analyzed using publicly available single-cell RNA-seq datasets. Astrocytes were identified by reference mapping with the Azimuth framework, and “pseudobulk” astrocyte expression profiles were generated by aggregating counts from astrocyte-annotated cells in each library. Differential expression between human and macaque astrocytes was then performed using *DESeq2*. Genes showing consistent species-biased expression in both comparisons were defined as evolutionarily affected genes (EAGs). Genes more highly expressed in human astrocytes were classified as human-enriched, whereas genes preferentially expressed in chimpanzee and macaque astrocytes were classified as non-human primate (NHP)-enriched. The volcano plot illustrates the relationship between the *DESeq*-derived *P*-value and LFC in human versus chimpanzee iAstrocytes in co-culture experiments; EAGs in primate astrocytes are highlighted in black and magenta. **e** Heatmap showing expression of selected EAGs in human and macaque fetal brain astrocytes. The expression of EAGs related to extracellular matrix organization, Hippo signaling genes are activated in the human lineage. Lipid handling, metabolic support, neuronal ion homeostasis, and synaptogenic signaling are significantly more expressed in NHP astrocytes than their human counterparts, is displayed (*DESeq2* method, genes defined in the analysis displayed in panel A are shown).

We observed robust and consistent differences in neuronal gene expression depending on the iAstrocyte species present in the co-culture (**Figure 2a**; *P*-adj. < 0.1 and |LFC| > 0.59, *DESeq2* method). Gene ontology analysis revealed enrichment of processes related to synaptic signaling, learning and memory, and cognition among genes upregulated in neurons exposed to chimpanzee iAstrocytes (**Figure 2b**). Consistent with this, marker gene analysis revealed coordinated upregulation of synaptic and activity-dependent programs in neurons co-cultured with chimpanzee iAstrocytes (**Supplementary Figure 6a** for the full gene set). These include genes involved in excitatory synaptic transmission (*Gria1*, *Grin2b*), synaptic scaffolding (*Dlg4/PSD95*, *Shank1*), vesicle trafficking, and activity-dependent signaling (*Camk2a*). Furthermore, loci associated with neuronal maturation, neuronal projection development and metabolism were also upregulated (**Supplementary Figure 6a**), consistent with a coordinated shift in neuronal maturation state rather than isolated pathway-specific effects.

In contrast, neurons exposed to human iAstrocytes upregulated genes associated with earlier developmental programs, including regulators of neuronal differentiation and axon guidance such as *Dcx*, *Draxin*, *Sema4c*, and *Nrg1*. Human iAstrocytes also induced upregulation of genes involved in RNA metabolism, including RNA-binding proteins and splicing factors (Hnrnp and Srsf families), as well as genes implicated in post-transcriptional regulation (**Figure 2b**). This transcriptional signature suggests that human astrocytes maintain neurons in a less mature, transcriptionally dynamic state, consistent with delayed functional maturation.

Given the prominent changes in genes involved in RNA metabolism, we next asked whether astrocyte species differentially regulate RNA processing in neurons. To test this, we performed differential exon usage analysis (*DEXSeq*).^71^ We observed increased exon inclusion in neurons exposed to human iAstrocytes compared to chimpanzee iAstrocytes (*P*-adj. < 0.05, |LFC| > 1; **Supplementary Figure 6b-c**). These changes affected genes involved in synaptic function and vesicle trafficking (**Supplementary Figure 6d**), revealing that species-specific differences in astrocyte-dependent RNA processing affect pathways associated with neuronal maturation. Together, these findings establish that human and chimpanzee iAstrocytes impose distinct transcriptional and post-transcriptional programs on neurons.

### Neuronal genes regulated by astrocytes are linked to neuropsychiatric disorders

The timing of neuronal maturation is a critical determinant of circuit function, and its disruption (heterochrony) is a recurrent feature of neurodevelopmental disorders, including autism, schizophrenia, and intellectual disability.^11,33,72–77^ Many neuronal genes that are preferentially induced by chimpanzee iAstrocytes, including *SHANK3*, *DLG4* (*PSD95*), *CAMK2A*, *SNAP25*, *GRIA1/3*, *GRIN1/2B*, and *GAD1* **(Supplementary Figure 6a**), are central to synaptic maturation and circuit stabilization. These genes are commonly disrupted in various neuropsychiatric disorders.

To determine whether these transcriptional changes are enriched in disease-relevant gene networks, we performed a disease-association analysis of genes upregulated in neurons exposed to chimpanzee compared to human iAstrocytes. This analysis revealed significant enrichment for gene sets associated with neurological and neuropsychiatric disorders, including schizophrenia, Alzheimer’s disease, dementia, and epilepsy (**Figure 2c**; Fisher’s test, Benjamini-Hochberg-corrected *P* < 0.01). In contrast, no significant enrichment was observed for genes upregulated in neurons co-cultured with human iAstrocytes.

These results indicate that astrocyte-dependent differences in neuronal maturation preferentially engage gene networks associated with brain disorders. Our results suggest a link between evolutionary divergence in astrocyte function and disease-relevant neuronal gene regulation.

### Evolutionary Changes in Astrocyte Gene Expression Programs

Astrocytes actively promote the formation and maturation of synapses,^38–40,78,79^ synaptic efficacy and plasticity, thereby shaping neuronal activity and network function.^80,81^ To identify the astrocyte-intrinsic molecular programs underlying the differential effects of human and chimpanzee iAstrocytes on neuronal maturation, we compared their transcriptomes in co-culture with rat neurons. To validate these results *in vivo*, we integrated single-cell RNA-seq profiles of human^82^ and macaque fetal brains.^83^ We focused only on genes that were consistently differentially expressed between human and non-human primate (NHP) astrocytes across both co-culture and *in vivo* conditions (**Methods;** Evolutionarily Affected Genes – EAGs; in both comparisons, *P*-adj. < 0.1, |LFC| > 0.59; *DESeq2* method; **Figure 2d**).

The analyses of EAGs revealed that human and NHP astrocytes activate distinct molecular programs when exposed to neurons. Human astrocytes showed a significant upregulation of genes involved in the formation of the extracellular matrix (ECM) and Hippo-TEAD mechanosensing pathway (**Figure 2e**, Fold Enrichment = 3, FDR = 0.06, Hypergeometric test). This transcriptional signature suggests that human astrocytes adopt an ECM-enriched, mechanosensing-regulated state relative to NHP astrocytes.

The human-biased EAG list also included primate-specific gene duplications such as NBPF family members and *SRGAP2B* (not shown). The coordinated upregulation of these loci suggests that evolutionary gene duplication events are embedded within this human-specific astrocytic regulatory architecture.

Notably, perhaps surprisingly, classical factors that regulate synaptic maturation, including glypicans,^39,42,79^ were not differentially expressed between human and NHP astrocytes (not shown). Furthermore, consistent with the observation that human and chimpanzee iAstrocytes do not differ in their capacity to induce structural synapses formation, thrombospondins 1 and 2 did not show differential expression in the inter-species comparisons (not shown). In addition, human astrocytes did not exhibit a broadly altered maturation score relative to NHPs in the presence of neurons (**Supplementary Figure 5c**), suggesting that the observed functional differences are not driven by global changes in astrocyte maturation but rather by selective regulation of particular neuron-maturation-supportive genes.

To identify these factors, we focused on genes upregulated in NHP astrocytes compared with human astrocytes. We found that both chimpanzee and macaque astrocytes upregulated a set of genes reflecting neuron-supportive astrocyte function. These included the core glutamate-handling and metabolic genes (*GLUL*, *PHGDH*) and key ion homeostasis factors (e.g., *ATP1A2*, *KCNK1*), consistent with the increased support for sustained neuronal activity of the NHP astrocytes relative to human cells (**Figure 2e**). Notably, astrocyte-microglial signaling factors (*SPARCL1*, *IL33*, *CX3CL1*, **Figure 2e**) were also enhanced in the NHP astrocytes. We interpret this result as a manifestation of a delay in synaptic pruning events in the human developing brain.

Notably, NHP astrocytes exhibited higher expression of genes involved in lipid handling than both iPSC and brain-derived human astrocytes. This includes the gene encoding apolipoprotein E (*APOE*) (**Figure 2e**, *P* < 0.01, *DESeq2* method), which was paralleled by an increased expression of APOE receptor *Lrp1* in rat neurons exposed to chimpanzee iAstrocytes (**Supplementary Figure 6d**), suggesting that brain evolution targets the astrocytic APOE – neuronal coupling. Crucially, *APOE* is the primary astrocyte-derived apolipoprotein involved in lipid transport to neurons and is associated with the regulation of synaptic function.^78,84–86^ Loss of APOE-dependent lipid delivery delays synaptic maturation,^78,85,87^ leading to higher-level brain dysfunctions.^88–90^ Therefore, given the selective effects on functional, rather than structural, synapse development observed in our system, we examined whether reduced astrocytic *APOE* expression contributes to diminished neuronal maturation in co-cultures with human astrocytes.

### Rescue of Neuronal Maturation Deficits by APOE Overexpression in Human iAstrocytes

Consistent with the transcriptomic data (**Figure 2e**), human iAstrocytes secreted significantly less APOE protein than chimpanzee iAstrocytes, both when cultured alone and in the presence of neurons (**Figure 3ab**; *P* < 0.01, *ANOVA*). In contrast, total soluble cholesterol levels in the culture medium were comparable between conditions (**Figure 3cd**), suggesting that reduced APOE secretion does not simply reflect differences in overall cholesterol availability. The dissociation between the overall APOE level and cholesterol content in the milieu suggests that APOE may regulate neuronal maturation through mechanisms beyond its role as cholesterol carrier. Therefore, we tested whether increasing APOE protein production in human iAstrocytes is sufficient to modulate neuronal activity in co-culture.

**Figure 3.**
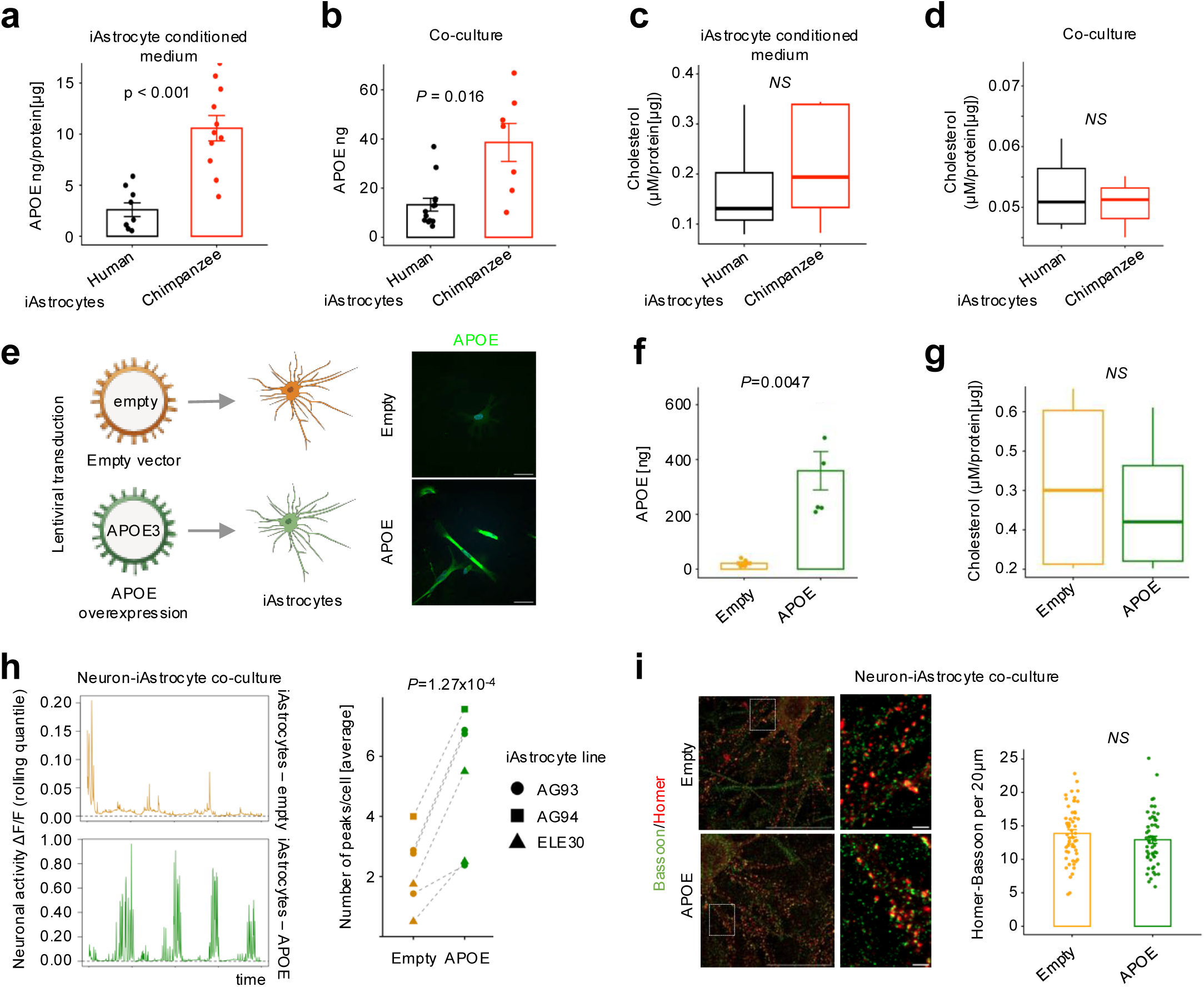
Reduced APOE secretion in human iAstrocytes delays neuronal maturation and can be rescued by APOE3 overexpression. **a** Quantification of APOE secretion by ELISA in conditioned media from human and chimpanzee iAstrocytes. Chimpanzee iAstrocytes secrete significantly higher levels of APOE protein. N experiments = 4. **b** APOE secretion measured in rat neuron-primate iAstrocyte co-cultures confirms higher APOE levels in chimpanzee astrocyte conditions. N experiments = 4. **c** Total soluble cholesterol levels measured in conditioned media from human and chimpanzee iAstrocyte cultures show no significant differences between species. N experiments = 3. **d** Total soluble cholesterol levels in rat-neuron human or chimpanzee iAstrocyte co-culture conditions remain comparable across conditions. N experiments = 3. **e** Lentiviral overexpression of APOE in human iAstrocytes. Left: overview of the experiment, right: representative images of immunofluorescence-based detection of APOE in cells transduced either with an empty vector or a vector coding APOE3. **f** Quantification of APOE secretion by ELISA in conditioned media from human iAstrocytes transduced with an empty vector or a vector expressing APOE3. N experiments = 3. **g** Total cholesterol levels in conditioned media from APOE-overexpressing iAstrocytes remain unchanged relative to control conditions (iAstrocytes transduced with an empty vector). N experiments = 3. **h** Quantification of neuronal activity showing increased frequency of calcium events in neurons co-cultured with APOE-overexpressing astrocytes. Rat hippocampal neurons were co-cultured with human iAstrocytes transduced with empty vector or APOE-expressing lentivirus. Spontaneous neuronal activity was assessed at DIV14 using GCaMP6f-based calcium imaging. Representative calcium traces from neurons co-cultured with control or APOE-expressing astrocytes are shown on the left. Quantification of peak frequency per cell across iAstrocyte lines is shown on the right. Each line represents an independent experiment. APOE significantly increased the frequency of calcium events (negative binomial mixed-effects model, *p* = 1.27 x 10^-4^). *N* = 5 experiments; 3 independent iAstrocyte lines. **i** Immunofluorescence analysis of excitatory synapses using Bassoon and Homer1 staining shows no change in total synapse number upon APOE overexpression, indicating that APOE regulates synaptic maturation rather than synapse formation. Each dot represents co-localizations along 20 μm of neuronal projection (N experiments = 3). Statistical differences between conditions were assessed using Student’s t-test. Scale bars in all images = 50 μm; insets: 2 μm.

Unlike most mammals, which carry a single APOE form, humans carry three common APOE isoforms (APOE2, APOE3, and APOE4) that differ in structure and lipid-binding properties. Chimpanzee APOE is the most similar to human APOE4,^91,92^ whereas APOE3 is the predominant isoform in modern human populations. Hence, to test whether APOE abundance is limiting instead of introducing an isoform comparison, we overexpressed human APOE3 in human iAstrocytes (**Figure 3e**) using lentiviral vectors. We observed a robust increase in *APOE* mRNA and protein secretion (**Figure 3ef**, *P* < 0.01, two-sided t-test) in human iAstrocytes. Importantly, APOE overexpression did not alter total soluble cholesterol levels in the culture medium (**Figure 3g**), recapitulating the dissociation between APOE abundance and bulk cholesterol levels observed between human and chimpanzee iAstrocytes.

Strikingly, neurons co-cultured with APOE-overexpressing iAstrocytes exhibited a significantly higher frequency of spontaneous calcium events than neurons co-cultured with empty vector-transduced human iAstrocytes. The increase corresponded to an approximately 2.1-fold increase in peak rate (**Figure 3h**, negative binomial mixed-effects model, β = 0.74 ± 0.19SE, *P* = 1.3x10^-4^). Notably, immunofluorescence analysis revealed no change in the density of structural synapses, marked by Bassoon-Homer colocalization (**Figure 3i**) when comparing neurons in co-cultures with iAstrocytes transduced with an empty versus APOE3 expression vector.

Together, these results demonstrate that increased astrocytic APOE is sufficient to promote functional maturation of excitatory synapses without altering synapse density. Consistent with the interspecies co-culture phenotypes, these findings identify APOE as a mechanistic contributor to differences in synaptic maturation between human and non-human primate astrocytes.

### Evolutionary Changes in DNA Regulatory Elements and their contacts at the APOE locus

Evolutionary changes in gene expression are frequently mediated by alterations in cis-regulatory elements and their activity.^93–97^ To investigate the regulatory basis of reduced APOE expression in human astrocytes, we examined chromatin accessibility and enhancer activity across the *APOE* locus in primate iAstrocytes.

Remarkably, comparative ATAC-seq and H3K27ac profiling^63^ revealed that the APOE promoter exhibits higher chromatin accessibility and enhancer-associated histone acetylation in NHP iAstrocytes compared to human iAstrocytes (**Figure 4a**). Chromatin conformation analysis^63^ (in situ Hi-C) showed that, despite overall preservation of domain organization, the APOE promoter forms a pronounced loop with a distal downstream regulatory element in chimpanzee iAstrocytes that is substantially weakened in human cells (**Figure 4a**). These observations indicate reduced promoter-enhancer dialogue at the *APOE* locus in human compared to NHP iAstrocytes.

**Figure 4.**
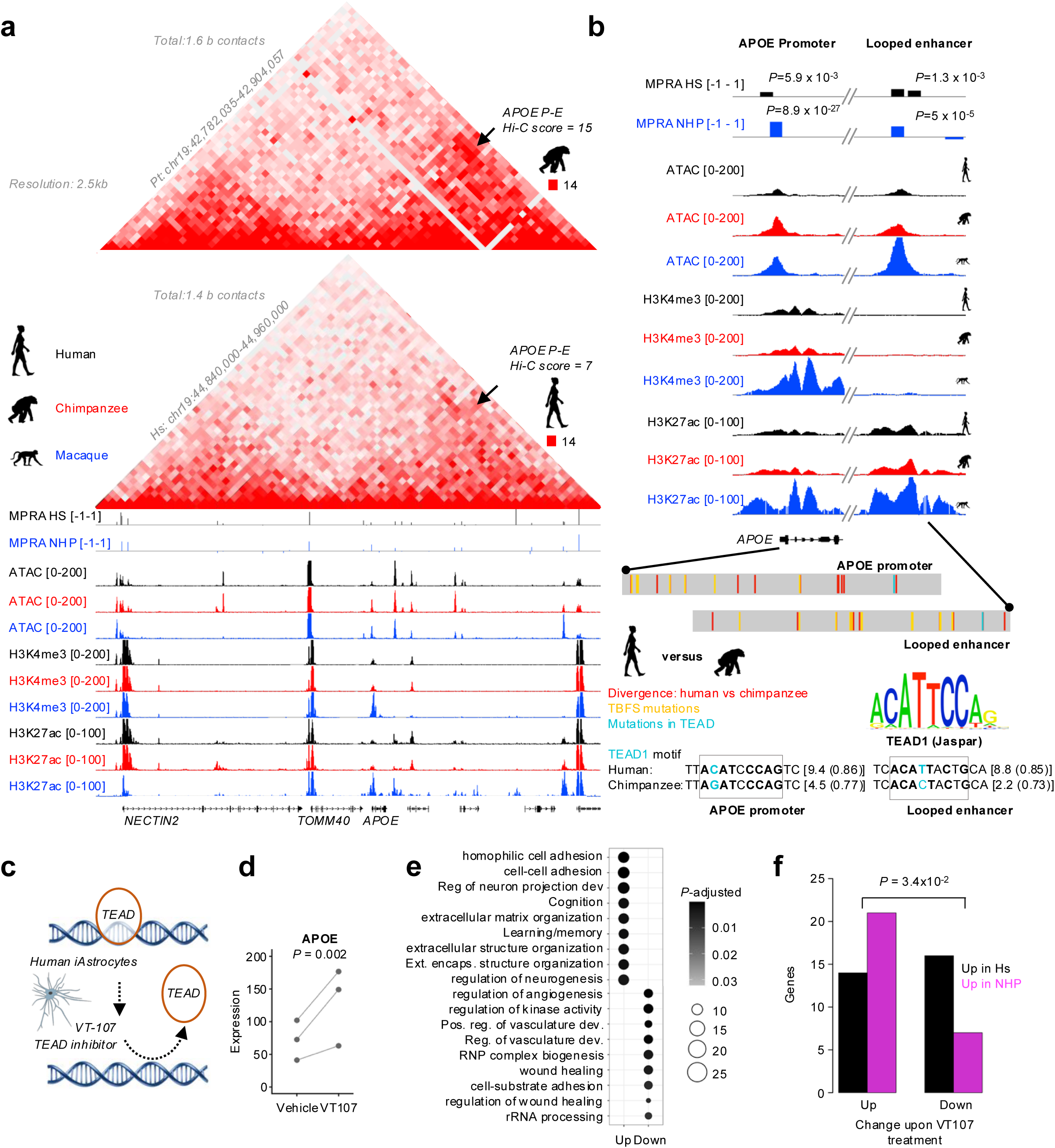
Evolutionary genomics of the APOE locus identifies Hippo-TEAD signaling as the contributor to the human-specific astrocyte transcriptional state. **a** A detailed view of regulatory elements at the APOE promoter and a distal enhancer engaged in chromatin looping. Comparative chromatin profiles reveal increased accessibility and enhancer activity in non-human primates. Right: schematic overview of the massively parallel reporter assay (MPRA) used to interrogate regulatory elements across the APOE locus. Candidate cis-regulatory elements derived from human, chimpanzee, and macaque genomes were cloned upstream of a minimal promoter and barcode reporter construct and transfected into astrocytoma cells. Regulatory activity of each sequence was quantified as the RNA/DNA barcode ratio. Panel displayed an integrative view of chromatin accessibility, histone modifications, and MPRA activity across the APOE locus in primate astrocytes. Hi-C data in human iAstrocytes reveal long-range chromatin interactions connecting the APOE promoter with distal regulatory regions. Tracks show MPRA activity together with ATAC-seq and histone modification profiles (H3K4me3 and H3K27ac) across human, chimpanzee, and macaque iAstrocytes. **b** Zoom in on chromatin modification and openness analysis at the APOE promoter and enhancer that interacts with the promoter via a chromatin loop. Bottom: evolutionary divergence in transcription factor binding sites within regulatory regions of the APOE locus. Human and chimpanzee orthologous sequences encompassing the APOE promoter and a distal looped enhancer were globally aligned. Red ticks indicate nucleotide differences between species that do not intersect TFBS. Yellow ticks mark evolutionary sequence changes that intersect predicted transcription factor binding sites (TFBS) identified in the human sequence using JASPAR position weight matrices. Cyan ticks indicate sites where sequence divergence alters predicted TEAD-family binding. For each TFBS detected in the human sequence, the orthologous chimpanzee sequence was extracted from the alignment and rescored with the same PWM. Bars show the difference in predicted binding affinity between species (Δ PWM score = human - chimpanzee). Analysis was restricted to high-confidence sites (normalized PWM score >0.84 in either species) for transcription factors expressed in human fetal brain astrocytes (among the top 25% of expressed genes in 30% of individuals). Sequence substitutions within TEAD1 motifs in both the promoter and enhancer increase the predicted TEAD binding score in the human lineage. The TEAD1 consensus motif derived from JASPAR is shown in the center. **c** Schematic representation of the inhibition of TEAD transcriptional activity using the auto-palmitoylation inhibitor VT-107. **d** APOE expression in human iAstrocytes is significantly increased upon TEAD inhibition (*P*–val.: *DESeq2* method. **e** Gene ontology enrichment analysis of genes differentially regulated upon TEAD inhibition (*DESeq2* method, *P*-adj.<0.1, |LFC| > 0.59, n=3 iAstrocyte lines). Genes upregulated following VT-107 treatment are enriched for processes related to neuronal development, extracellular matrix organization, and cognitive functions. **f** Comparison of TEAD-responsive genes with evolutionarily affected genes in astrocytes. Genes upregulated upon TEAD inhibition are significantly enriched among genes that show higher expression in non-human primate astrocytes relative to human astrocytes.

To functionally interrogate regulatory elements across the *APOE* locus, we performed a massively parallel reporter assay (MPRA, Duński et al., in preparation) using candidate sequences selected from integrated ATAC-seq and chromatin interaction data spanning ±500 kb around the *APOE* promoter. Reporter assays (**Methods**) revealed numerous active regulatory elements (**Figure 4a**, **Supplementary Figure 7a-d**), with most human sequences exhibiting comparable or higher activity than their NHP counterparts (**Supplementary Figure 7e**; *P* = 0.049, two-sided t-test), indicating a marked regulatory divergence at the locus.

Interestingly, the *APOE* promoter itself was a notable exception: the chimpanzee promoter sequence consistently drove higher reporter expression than its human ortholog (**Figure 4b**). The distal enhancer, that physically contacts the *APOE* promoter in the NHP iAstrocytes, displayed a similar pattern, with stronger reporter activity, chromatin accessibility, and H3K27ac enrichment in NHP cells. Hence, reduced *APOE* expression in human astrocytes is associated with locus-specific changes affecting both promoter and enhancer activity and loop formation.

Next, we sought to identify sequence differences within *APOE* regulatory elements that could explain the evolutionary changes in APOE expression. Comparative sequence analysis revealed species-specific differences in transcription factor binding potential across *APOE* promoter and enhancer elements (**Figure 4b**, **Supplementary Figure 7a-d**). Motifs for several astrocyte-associated transcription factors showed higher predicted affinity in chimpanzee sequences (**Supplementary Figure 7f**), consistent with increased *APOE* mRNA production in the NHPs. In contrast, motifs recognized by TEAD transcription factors showed stronger concordance with the consensus sequence in human regulatory elements at both the *APOE* promoter and distal enhancer, identifying TEAD as a candidate regulator of species-specific APOE expression. We decided to test this hypothesis.

### Hippo-TEAD Signaling Links Astrocyte Evolution to Delayed Maturation Programs

To assess whether TEAD activity regulates species-specific astrocyte transcriptional programs, including *APOE* expression, we inhibited TEAD signaling in human iAstrocytes using VT-107, a compound that blocks TEAD auto-palmitoylation, thereby blocking TEAD-mediated transcriptional programs (iAstrocyte lines from 3 independent individuals; **Figure 4c**).

RNA-seq analysis revealed a coordinated suppression of canonical YAP/TAZ-TEAD target genes (|LFC|>0.59, *P*-adj. < 0.1; *DESeq2* method; **Supplementary Figure 8a**), including *CCN1*, *CCN2*, and *ANKRD1*. This was accompanied by downregulation of key regulators of proliferative programs (*MYC*, *CCND1*) and genes associated with growth factor signaling and migratory states (e.g., *TGFB2* and *AXL*), confirming effective attenuation of TEAD activity.

Notably, we found that TEAD inhibition resulted in a robust upregulation of *APOE* expression in human iAstrocytes (**Figure 4d**; *P* = 0.002, *DESeq2* method). Hence, evolutionary enhancement of the TEAD recognition motif, both within the *APOE* promoter and in its looped enhancer, likely leads to diminished *APOE* expression in the human lineage.

We next decided to address the effect of TEAD inhibition on astrocyte transcriptome more broadly. GO term analysis revealed that TEAD block induced a transcriptional program enriched for neuronal development and synaptic function in human iAstrocytes (**Figure 4e**; **Supplementary Figure 8b**). Upregulated genes included key mediators of neuron-glia interaction and synaptic organization, such as *NRXN3 and NLGN3*, as well as axon guidance molecules (e.g., *SEMA3E and EFNB3*) and genes involved in cell-cell adhesion, including multiple protocadherins, consistent with enhanced capacity for intercellular communication upon TEAD block.

The pathways we observed were indicative of a more neuronal-maturation promoting program elicited by TEAD inhibition in the human iAstrocytes (**Supplementary Figure 8b**), which suggested that TEAD activation in the human lineage might be responsible to some of the aspects of the astrocyte evolutionary divergence in primates. To test this possibility, we compared TEAD-responsive genes with species-biased expression programs in human and NHP astrocytes. We found that genes induced upon TEAD inhibition significantly overlapped with the transcripts enriched in NHP astrocytes, whereas genes suppressed by the VT-107 treatment were enriched among those upregulated in the human cells (**Figure 4f**; *P* = 3.4 x 10^-2^, Fisher’s exact test). Thus, sustained TEAD activity significantly contributes to the evolution of human-specific astrocyte gene expression program. Likewise, these results suggest the role of mechanosensing in the regulation of neuron-supportive features of astrocytes (**Figure 5**).

**Figure 5.**
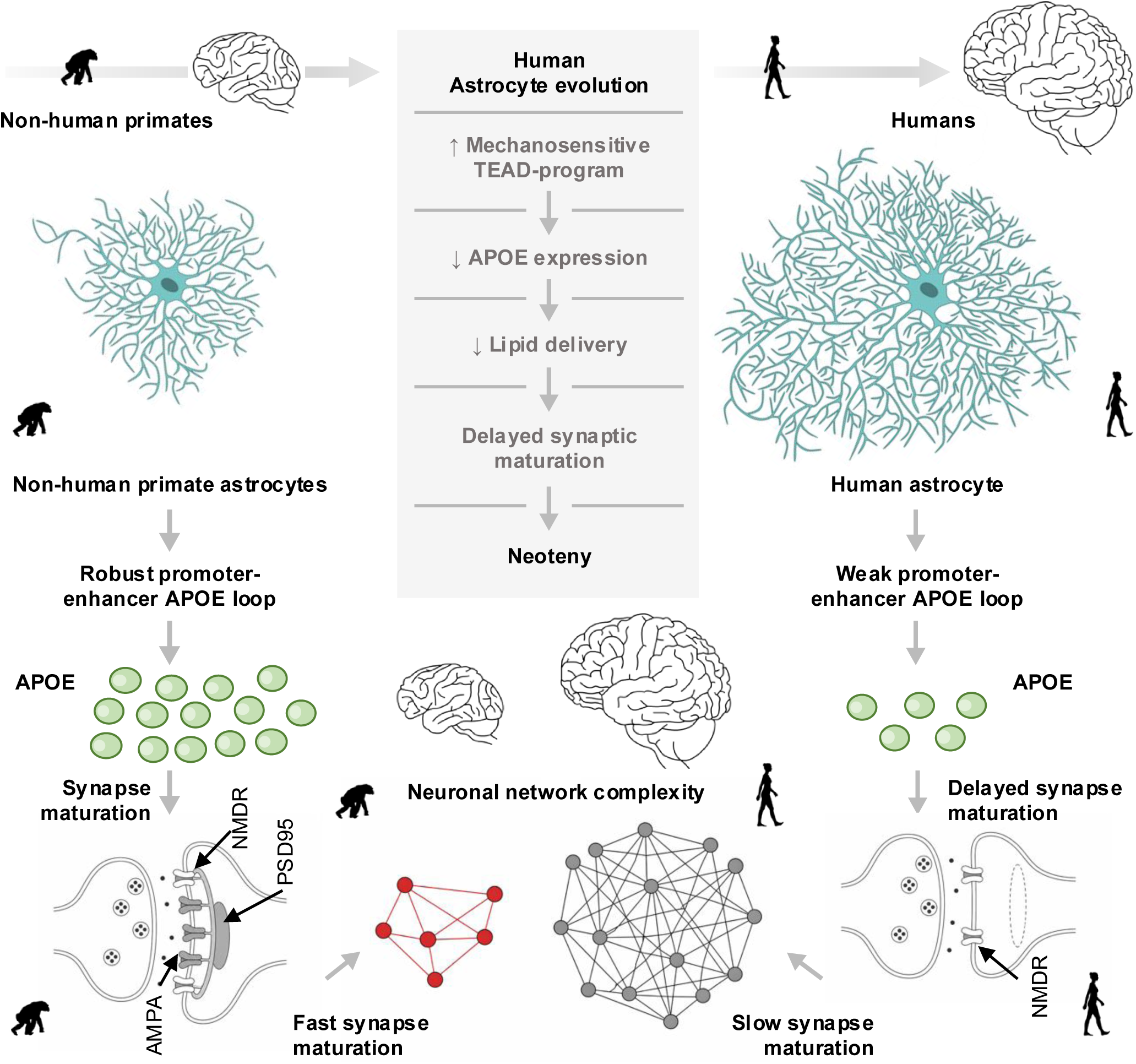
Model: evolutionary remodeling of astrocyte APOE support delays synapse maturation and shapes neuronal network development in humans. Our model shows how evolutionary changes in astrocyte transcriptional programs influence neuronal maturation and circuit architecture. Non-human primate astrocytes exhibit strong APOE promoter-enhancer loop formation, reflected by high APOE secretion. This promotes rapid synaptic maturation, characterized by recruitment of AMPA receptors to initially NMDA-only (“silent”) synapses, resulting in early stabilization of neuronal circuits. In contrast, human astrocytes display an evolutionarily reinforced Hippo-TEAD transcriptional program that represses APOE expression and reduces astrocyte-derived lipid delivery. Limited lipid support delays AMPA receptor recruitment, prolonging the presence of NMDA-only synapses and slowing synaptic maturation. Network schematics illustrate the resulting differences in circuit dynamics: rapid synapse maturation promotes early stabilization of relatively simple networks, whereas delayed maturation extends circuit plasticity and permits the emergence of more complex neuronal connectivity. Hence, this Supplementary developmental window maintains circuits in a plastic state and contributes to the prolonged neuronal developmental trajectory (neoteny) characteristic of the human brain.

Taken together, we identify astrocytes as active regulators of the timing of neuronal maturation, with their role having evolved in the human lineage (**Figure 5**). We show that increased Hippo-TEAD expression in human astrocytes, likely tightly linked with their expanded size and complexity, represses neuron-supportive and lipid metabolic programs, including robust APOE production. This in turn, delays neuronal maturation. Attenuation of TEAD signaling induces neuron-supportive transcriptional programs observed in non-human primate astrocytes. Hence, evolutionary changes in astrocyte gene regulation shape neuronal maturation, linking astroglial evolution to species-specific differences in the timing of brain development.

## Discussion

Neuronal maturation in mammals follows a conserved sequence of molecular and functional transitions, yet its tempo varies substantially across species and is markedly prolonged in humans. These differences have been largely attributed to neuron-intrinsic phenomena,^28,29,98^ and entail variation in biochemical reaction kinetics, protein stability, and metabolic state,^22–27,95^ suggesting that developmental time is encoded within neurons.^22–27,95^ Our findings overturn this model. We demonstrate that neuronal maturation tempo is actively imposed by astrocytes, and that this non-cell-autonomous control has been fundamentally rewired during human evolution. Using genetically identical neurons as a neutral substrate, we show that human and chimpanzee astrocytes, derived from species that diverged fewer than seven million years ago, shape neuronal developmental trajectories in an opposing way: chimpanzee astrocytes accelerate maturation, while human astrocytes delay it. Hence, brain neoteny is, in part, a glial phenomenon.

Astrocytes have undergone substantial evolutionary changes in mammals, including increases in size and complexity and marked changes in gene expression programs in humans relative to other species.^54–63^ The relevance of these evolutionary innovations remains, however, unclear. It is now well recognized that rodent astrocytes promote functional maturation of human iPSC-derived neurons more effectively than human astrocytes.^99^ This phenomenon was previously attributed to technical limitations of iPSC protocols. Our data suggest, that it instead reflects a conserved feature of human astrocyte biology. Previous, groundbreaking study revealed that human astrocytes enhance synaptic plasticity and improve learning when transplanted into mouse brains,^56^ demonstrating that astrocyte evolution contributed to enhanced cognitive abilities. Our findings resonate with both observations and lead to a hypothesis, that by slowing down developmental trajectory of neurons, human astrocytes expand neuronal plasticity and contribute to the formation of a more complex connectome. This proposal will need to be critically evaluated in chimera systems.

A key question in evolutionary neuroscience is whether human brain development represents a simple temporal extension of the ancestral primate program or if it involves a qualitative reorganization of developmental processes. Single-cell transcriptomics of the developing human brain revealed a complex answer to this question. Distinct progenitor populations and areal specification programs arise with greater intricacy in humans than in other primates, suggesting that human brain evolution involves both quantitative and qualitative changes.^100–102^ Our findings extend this view and show that human astrocytes do not simply provide developmental support at a slower pace, but deploy a fundamentally distinct regulatory program that actively imposes delayed neuronal maturation.

APOE is best known for its role in lipid metabolism and as a major genetic risk factor for Alzheimer’s disease.^103–107^ The APOE gene exhibits signatures of positive selection in humans,^108^ raising the possibility that changes in its regulation were adaptive. APOE is polymorphic in humans and exists as three isoforms (APOE2, APOE3, and APOE4), whereas rodents and NHP possess a single isoform reminiscent of the APOE4 form.^92,109^ While it is unclear which evolutionary forces shape APOE genetics, it has been recognized that its isoforms play important roles in neurodevelopment, including synapse formation, synaptic plasticity, and neuronal repair.^110–119^ Notably, the APOE4 variant is associated with reduced astrocytic APOE secretion, impaired lipid transport, and reduced synaptic support relative to APOE2 or APOE3^110–119^ phenotypes that parallel the reduced pro-maturational capacity of human astrocytes we describe here. However, APOE4-carrying human neurons themselves exhibit enhanced synaptogenesis relative to APOE3 neurons,^120^ suggesting that APOE exerts distinct, cell-type-specific effects on synaptic development in neurons and astrocytes. Our data reveal that chimpanzee iAstrocytes, which express an APOE4-like isoform, strongly promote synaptic maturation. This is inconsistent with models in which isoform-specific properties alone determine synaptic outcomes and instead points to APOE dosage as the dominant regulatory variable. The sufficiency of increased APOE3 expression in human iAstrocytes to accelerate neuronal maturation further supports the conclusion that total astrocyte-derived APOE output, rather than isoform identity, is the primary determinant of its role in developmental timing. In line with this proposal, snRNA-seq profiling human brain development^121^ reveals that APOE expression in astrocytes is low during fetal stages and increases markedly after birth. Thus APOE expression aligns with the postnatal emergence of astrocyte-dependent neuronal maturation program described here.

Likewise, the emergence of APOE3 in humans approximately 200,000 years ago^91,109^ has puzzled evolutionary biologists. Dietary adaptation and protection against age-related disease might be the root cause of the emergence of APOE3 and APOE2 isoforms in the human lineage.^108,122^ Finch and Sapolsky proposed that evolutionary selection favored parents who remained cognitively intact for longer because human offspring mature exceptionally slowly and require years of parental care, thereby creating a pressure against early-onset neurodegeneration and for the neuroprotective APOE3 allele. Our findings suggest that APOE may be upstream of this evolutionary logic: if the dosage of astrocyte-derived APOE shapes the tempo of neuronal maturation, then changes in APOE regulation may have contributed to creating the very prolonged developmental trajectory that necessitated Supplementary parental care in the first place. In this view, Altogehter, in this view, APOE evolution shaped not only who could survive long enough to raise slowly-developing children, but how slowly those children developed.

APOE may influence neuronal maturation through both metabolic and signaling functions. In addition to supplying lipids required for dendritic spine growth and synaptic membrane biogenesis, APOE engages LDL receptor family members, including LRP1, to regulate postsynaptic membrane composition, AMPAR trafficking, and NMDAR-associated signaling pathways central to synapse maturation.^103,123^ Our data show that *LRP1* is upregulated in rat neurons in the presence of chimpanzee iAstrocytes, indicating a positive feedback loop. Yet, this observation does not rule out the possibility that APOE lipoproteins are composed of different molecules in human and NHP iAstrocytes, and that their identity matters for the timing of neuronal maturation. Recently, APOE-containing lipoprotein particles have been reported to carry regulatory molecules, including microRNAs, that can influence neuronal gene expression and metabolic programs.^124^ Hence, APOE levels might modulate developmental tempo through a combination of several pathways. In future work, it will be important to dissect how APOE particles impact synapse maturation.

The Hippo-TEAD^130,131^ pathway is a highly conserved cascade regulating organ size, cell proliferation, and development of the nervous system.^125,126^ Because YAP/TAZ-TEAD signaling integrates mechanical cues with transcriptional control, it provides a plausible molecular framework linking the remarkable morphological expansion of human astrocytes to their evolutionarily acquired functions. Here, we identify APOE as a downstream target of Hippo-TEAD signaling, suggesting that this pathway links astrocyte morphology to neuronal developmental timing.

Our data further suggest that this regulatory axis may be embedded within a broader remodeling of cellular programs involved in sensing and responding to the extracellular mechanical environment. Human astrocytes upregulate a coherent set of mechanosensing-associated genes, including *YAP1* and *PIEZO1*. Notably, *PIEZO1* is selectively enriched in glia rather than in neurons in the human cerebral cortex (https://scapex.ethz.ch/; scRNA-seq data from ref.^61^, accessed 28.06.2026), raising the possibility that the evolutionary expansion of human astrocytes was accompanied by an increased capacity to sense and respond to extracellular mechanical cues. This hypothesis remains to be tested experimentally.

This mechanosensory remodeling may also carry a cost in terms of disease: ECM dysregulation is among the most robust molecular signatures of schizophrenia, with disease-related genetic variants mapping to ECM-encoding genes,^127–131^ while the prolonged developmental trajectory of the human brain increases the window of vulnerability to genetic perturbations.^13,20,21^ Consistent with this, genes differentially expressed in neurons depending on iAstrocyte species are enriched for loci linked to schizophrenia, ASD, and Alzheimer’s disease. This view is consistent with increasing evidence that astrocyte dysfunction contributes to neurodevelopmental, neuropsychiatric and neurodegenerative disorders.^37,132^ Together, our findings raise the possibility that astrocytes contribute to neuropsychiatric susceptibility by regulating the tempo of neuronal maturation and the duration of developmental plasticity windows. This would constitute a glial mechanism linking brain evolution to disease vulnerability.

Previous studies demonstrated that human iPSC-derived neurons mature more slowly than NHP counterparts even when supported by the same rodent astrocytes, providing compelling evidence that neuronal developmental tempo is at least partly encoded by neuron-intrinsic programs.^28,29,35,98^ In a companion study (Rava et al., *Biorxiv* 2026 Evolutionary diversification of lipid logistics shapes synaptic maturation in primates), Rava et al. further show that delayed maturation of human neurons is associated with altered neuronal lipid handling relative to chimpanzees. Our findings extend this model by identifying an independently evolved astrocytic mechanism that converges on the same biological process. Together, our two studies suggest that lipid homeostasis represents a central regulatory node controlling the pace of neuronal maturation, integrating both neuron-intrinsic metabolic programs and astrocyte-derived signals. Thus, neoteny emerges from the interplay between cell-intrinsic neuronal properties and non-cell-autonomous astrocyte signals: two separable and independently evolved layers of neuronal developmental control.

## Acknowledgements

Work in the Pękowska lab is funded by the Dioscuri Grant (Dioscuri is a program initiated by the Max Planck Society (MPG), jointly managed with the National Science Centre in Poland (NCN), and mutually funded by the Polish Ministry of Education and Science and the German Federal Ministry of Education and Research (BMBF)). Confocal imaging was performed at the Laboratory of Imaging Tissue Structure and Function, an imaging core facility at the Nencki Institute of Experimental Biology and part of the infrastructure of the Polish Euro-BioImaging Node. High performance computing was performed at Poznan Supercomputing and Networking Centre (https://www.pcss.pl/). We would like to acknowledge Piotr Michaluk from the Nencki Institute and the Municipal Zoological Garden in Warsaw, in particular Agnieszka Czujkowska and Andrzej Kruszewicz.

## Author contributions

Conceptualization and study design: A.Pe.; funding acquisition: A.Pe., E.D., and K.C.; experimental work: K.C., I.F., E.D., and B.D.; co-culture establishment and troubleshooting: I.F. and K.C.; analysis design and experimental analyses: K.C.; morphological analysis: G.V.; iPS cell generation and astrocyte culture: A.P.; astrocyte validation: K.H. and E.B.; bioinformatic analysis: A.Pe., with assistance from M.A. and B.H.; additional experimental support: V.R. and E.T.; supervision of co-culture system: J.W.; writing - original draft: A.Pe.

## Methods

### Cell lines

Human iPS cell line “ELE30” (IIMCBi002-A) was a gift from Prof. J. Jaworski and “GM3651”, “AG9319”, and “AG9429” iPS cell lines were a gift from Prof. F. Gage. Chimpanzee iPS cell lines “Mandy04” and “Mandy06” were generated in our laboratory as described previously (Ciuba et al., 2025).

The chimpanzee “Kimby12” iPS line was generated in our laboratory. Small bits of leftover skin tissue, from a scheduled medical procedure performed in the Warsaw ZOO, were obtained from 34 years-old chimpanzee “Kimberly” (female). The skin fragments were washed and cut into small pieces before being plated at 3-4 explants per well of a 6-well plate coated with 0.2% gelatin (Merck). The explants were then covered with 400 μl of culture medium (CM), which was a mixture of high-glucose DMEM (Biowest), 20% FBS (Merck) and 1% penicillin-streptomycin (Thermo Fisher). The CM was changed every other day until the explants had attached to the culture plate. The fibroblasts then migrated out of the explants and attached to the culture plate. After three weeks of culture, the fibroblasts were passaged using Trypsin-EDTA (Merck) and used for reprogramming in the first passage. 5,000 fibroblasts/cm² were seeded onto hESC-qualified Matrigel (Corning)-coated wells. Reprogramming was initiated on day 5 post-plating using the ReproRNA-OKSGM Kit (STEMCELL Technologies). Briefly, the conditioned medium (CM) was replaced with growth medium (GM) (Advanced DMEM (Thermo Fisher Scientific) supplemented with 10% FBS (Merck), 2 mM L-glutamine (Thermo Fisher Scientific) and 175 ng/ml recombinant B18R protein (STEMCELL Technologies)). The fibroblasts were transfected on day 0 with a freshly prepared ReproRNA cocktail (2 μl ReproRNA^(TM)-OKSGM (STEMCELL Technologies, 05931), 4 μl ReproRNA Transfection Supplement (STEMCELL Technologies) and 4 μl ReproRNA Transfection Reagent (STEMCELL Technologies) in 200 μl Opti-MEM Reduced-Serum Medium (Thermo Fisher). The medium was changed daily during days 1–7 post-transfection (GM supplemented with 0.4 ng/ml puromycin (Thermo Fisher)). Pluripotent stem cell selection was carried out between days 8 and 14 post-transfection by culturing the cells in ReproTeSR (STEMCELL Technologies), supplemented with 175 ng/ml B18R protein. From day 14 post-transfection, the cells were cultured in ReproTeSR without B18R protein until iPS cell-like colonies appeared (days 17–20 post-transfection). The iPS cell-like colonies were manually picked, placed in separate wells of a 24-well plate and dislodged using a pipette tip. The colonies were further expanded and used for downstream applications.

### Neural progenitor generation from iPSc cells

To initiate differentiation into iNP cells, iPS cell colonies were detached and dislodged into single cells using Accutase (Merck; Day 0). Three million iPS cells were then resuspended in 1 ml of STEMdiff Neural Induction Medium + SMADi (STEMCELL Technologies) and seeded into a single well of a 24-well AggreWell 800 plate in the presence of 10 μM Y-27632 (Adooq Bioscience) to form embryoid bodies (EBs). ¾ of the medium was replaced daily between days 1 and 4. On day 5, the EBs were collected using wide-bore tips and 37 µM reversible cell strainers (STEMCELL Technologies), then plated into a single well of a 6-well plate coated with hESc-qualified Matrigel in STEMdiff Neural Induction Medium + SMADi. A full medium change was performed daily from days 6 to 11. On day 12, the neural rosettes were selected using STEMdiff Neural Rosette Selection Reagent (STEMCELL Technologies) and replated into a single well of a six-well plate coated with Matrigel. The cells were cultured with a daily medium change until day 14 for chimpanzees, or day 19 for humans. A single-cell suspension of iNPs was then obtained using Accutase. The cells were plated in Matrigel-coated plates and cultured in STEMdiff Neural Progenitor Medium (STEMCELL Technologies), and passaged using Accutase in 1:5-1:5 ratio every five to six days.

### iAstrocyte differentiation and culture

iAstrocytes were obtained from a purified (CD271-/CD133+/CD184+) population of iNPs. A single-cell suspension of iNP cells was immunolabelled with antibodies against CD271 (BD Biosciences, dilution 1:20), CD133 (Miltenyi Biotec, dilution 1:100), and CD184 (BD Biosciences, dilution 1:5). The CD271-/CD133+/CD184+ iNPs were then purified using flow cytometry. iAstrocyte differentiation was induced between passages 2–5, following flow cytometry-assisted purification.

For differentiation, the iNPs were detached with Accutase (Merck) and dislodged to single cells by gently pipetting. 150,000 iNPs/cm^2^ were seeded on hESC-Matrigel (Corning) in iAstrocyte medium (ScienCell). On in vitro culture day 14 (DIV14), the cells were detched, with Accutase (Merck) and dislodged into a suspension of single cells. Following this, the cells were counted and seeded back at the original density in the respective vessels. On DIV28, the cells were detached again and seeded for validation procedures. During differentiation and culture, half of the cell media was replaced with fresh iAstrocyte medium every other day.

### Primary hippocampal cultures

Neuronal primary cultures were obtained from the brains of Wistar rats on postnatal day 0 (P0), which were sacrificed by decapitation. The dissected hippocampi were collected in an ice-cold DM-KYN/Mg solution containing 1 M Na2SO 4, 0.5 M K2SO 4, 1 M MgCl2, 0.1 M CaCl2, 1 M HEPES, 3 M glucose and 10 mM kynurenic acid. After washing in the same buffer solution, the tissue was incubated in DM-KYN/Mg containing 10 U/ml papain (Worthington Biochemicals), at 37°C for twenty minutes. Next, the tissue was rinsed three times in warm DM-KYN/Mg, and then incubated with a 10 gm/ml trypsin inhibitor (Merck) solution in DM-KYN/Mg, at 37°C for five minutes. The hippocampi were rinsed again in a neuronal plating medium containing Minimum Essential Medium (Thermo Fisher Scientific), 10% fetal bovine serum (Thermo Fisher Scientific), 1% MEM non-essential amino acid solution (Merck), 1 mM sodium pyruvate (Thermo Fisher Scientific), 3 mM glucose (Merck), 1x GlutaMAX (Thermo Fisher Scientific) and 100 μg/ml penicillin–streptomycin (1x P/S, Merck). The suspension was then gently triturated 20 times with a 1 ml pipette tip to mechanically dissociate the tissue. After centrifuging the suspension at 180 x g for 10 minutes, the supernatant was discarded, after which the pellet was resuspended in 1 ml of plating medium. The number of viable cells was evaluated using trypan blue in a hemocytometer. The cells were plated in neuronal plating media at a density of 100,000 cells per 13 mm diameter coverslip (VWR) which was coated with 50 µg/ml poly-D-lysine (Merck) and 5 µg/ml laminin (Merck). Once the cells had adhered (approximately one hour after plating), the plating medium was replaced with neural growth medium (NGM) containing Neurobasal (Thermo Fisher Scientific), B-27 supplement (1x, A3582801, Thermo Fisher Scientific), GlutaMAX supplement (1x) and P/S (1x). On the third day in vitro (DIV3), the respective cultures were treated with 75 µM 5-fluoro-2′-deoxyuridine (FdU; Merck) to inhibit glial proliferation, resulting in an almost pure neuronal culture. ½ of the medium was exchanged every 4th day.

### Co-cultures

All morphological analyses, synapse immunofluorescence stainings and GCAMP activity assays were performed using direct co-cultures. For viability assays, indirect co-cultures were used.

In brief, 50,000 iAstrocytes were plated 24 hours before the addition of neurons in Astrocyte Medium (ScienCell), onto hESC Matrigel-coated 13 mm diameter coverslips (VWR) for direct co-cultures, or onto transwell inserts (VWR) for indirect co-cultures. After 24 hours, the medium was exchanged for neuronal plating media, and 100,000 rat hippocampal neurons (isolated as described above) were plated on top of the iAstrocytes. Alternatively, neurons were plated to PDL/Laminin-coated coverslips and astrocyte-containing inserts were placed on top of them. The cultures were treated with 75 μM FdU (Merck) at DIV3. During further culture, ½ of the medium was exchanged every 4th day. The co-cultures were kept until DIV14, when they were used for the relevant experiments.

### LN-229 cell culture

LN-229 (ATCC catalogue numr CRL-2611) were roasnely cultured on cell culture treat-d plastic dishes (Corning). Cells were maintained in DMEM/F-12, GlutaMAX™ (Gibco) supplemented with 10% Fetal bovine serum (SIGMA). Cells were fed every 3 days. Cells were routinely passaged upon reaching confluence using accutase (Merck) and seeded onto fresh plates at density of 1:8 to 1:4

### Neuronal viability assay

Viability assays were performed on DIV14. Briefly, the medium was removed and the neurons were washed once with Dulbecco’s phosphate buffered saline with Ca2+ and Mg2+ (DPBS with ions; VWR). Then, staining was performed using the LIVE/DEAD Cytotoxicity Kit (Thermo Fisher Scientific), according to manufacturer’s instructions. After staining, the cells were rinsed once (DPBS with ions), mounted on the microscope slides using ProLong Diamond Antifade mountant (Thermo Fisher Scientific) and immediately imaged using ZOE Fluorescence Imager (Bio-Rad). Images of three random areas per coverslip for each condition were acquired in the green channel (Calcein, live cells) and red channel (PI, dead cells).

### Neuronal transfection

To visualize neuronal morphology and dendritic spines, the cultures were transfected with a plasmid vector expressing enhanced green fluorescent protein (eGFP) under the human synapsin (hSYN) promoter (gift prom Prof. J. Jaworski). To monitor spontaneous neuronal activity in live imaging, we used pZac-GCaMP6f vector under synapsin promoter (gift from Dr. P. Michaluk).

Transfections were performed at DIV 7 using Lipofectamine 2000 (Thermo Fisher Scientific) according to the manufacturer’s protocol. 0.5 μg of total DNA and 1μl of Lipofectamine 2000 were used per coverslip. Before adding the transfection mixture, half of the cell media was collected and stored in the respective tubes. The cells were then treated with the transfection mixture and incubated for one hour. The mixture was then removed and replaced with 200 μl of fresh medium containing B-27 supplement mixed with 200 μl of the previously collected medium. 1/3 of the medium was subsequently replaced twice a week.

### Morphometric analysis of dendritic spines

At DIV 14, the cultures were fixed with a mixture of 4% PFA (Merck) 4% sucrose (Merck) in DPBS for 10 minutes at room temperature (RT). Coverslips were rinsed three times with DPBS with ions and mounted on microscope slides using Fluoromount-G Mounting Medium (Thermo Fisher Scientific).

Images of neuronal dendrites were acquired using a Zeiss LSM780 confocal microscope with a 63×/1.4 NA oil immersion objective and a 488 nm laser set to 3.5% transmission, with a pixel count of 1024×1024. A series of z-stacks were collected for each cell with a step size of 0.4 μm, and additional digital zoom resulted in a lateral resolution of 0.07 μm per pixel. Morphometric analysis of the dendritic spines was performed semi-automatically using SpineMagick software (patent no. WO/2013/021001), as previously described.^133^ Spines of the secondary and tertiary dendrites were analyzed to reduce possible differences in spine morphology caused by their location on dendrites of different ranks. We used a scale-free parameter reflecting spine shape, namely the relative change in the spine length-to-head width ratio. Spine length was determined by measuring the curvilinear length along a fitted virtual skeleton of the spine. This procedure involved looking for a curve along which integrated fluorescence was at a maximum. The head width was defined as the diameter of the largest spine section, excluding the bottom part of the spine (one-third of the spine length adjacent to the dendrite). The dendritic spines of three primary hippocampal cultures were analyzed.

### Sholl Analysis of neurons

Sholl analysis was performed to assess the dendritic complexity of eGFP-Synapsin-transfected neurons in the presence and absence of FDU, as well as in neurons co-cultured with primate astrocytes. Images of the neurons were acquired using a Zeiss LSM780 confocal microscope fitted with a 40x/1.4 NA oil immersion objective and a 488 nm laser. The neuronal projections were manually traced using the NeuronJ 1.4.3 plugin in Fiji. Sholl analysis was then performed on the skeletons generated from these tracings using the Sholl Analysis v4.2.1 plugin in Fiji.^134^

### Live imaging of spontaneous neuronal activity

For live imaging of the spontaneous neuronal activity, the neurons were transfected with pZac-GCaMP-6f plasmid, as described above. Live imaging of the neurons was performed at DIV14 with Zeiss 7 Cell Discoverer microscope in LSM mode, using 20x/0.7 dry objective in 0.5x magnification. Images were acquired every 500 milliseconds for a total of 7 minutes per field of view. Region of interest (ROIs) for fluorescence analyses were manually selected within the cell body.

### Calcium Imaging Trace Visualization

Time-series fluorescence data were analyzed and visualized using R Studio (ver. 2025.05.1+513) with the *tidyverse* package collection. Raw fluorescence traces were imported as tab-delimited text files, reshaped from wide to long format, and organized by experiment, species, condition, and cell identity. To remove initial recording instability, the first 50 frames of each trace were excluded, and time was re-indexed starting from frame 1. For baseline correction, the 10th percentile of fluorescence intensity was calculated separately for each cell, and this value was subtracted from the corresponding trace. Fluorescence values below this threshold were set to zero to suppress low-intensity baseline fluctuations while preserving relative signal amplitude. Corrected traces were visualized as time-series plots without amplitude normalization. Figures were generated using the R/Bioconductor package *ggplot2*.

### Immunofluorescence

At DIV 14, the cultures were fixed with a mixture of 4% PFA (Merck) 4% sucrose (Merck) in DPBS (10 minutes, RT). All permeabilizing, blocking and washing solutions were prepared using DBBP with ions. They were then rinsed three times with DPBS and permeabilized (5 minutes, RT) with 0.5% Triton X-100 (Merck) in DPBS. The permeabilized cells were then washed three times for five minutes each with DPBS. The cells were then incubated in a blocking solution: 0.5% BSA (BioShop) in DPBS (1 hour, RT). The cells were then incubated with the respective primary antibodies, diluted in the blocking solution as required (1 hour, RT). After the primary antibody incubation, the cells were rinsed with DPBS three times for five minutes each and incubated with the respective AlexaFluor fluorescently labelled secondary antibodies (Thermo Fisher Scientific) and Hoechst 33342 (Thermo Fisher Scientific) for nuclear staining. After three further washes with DPBS, the cells were mounted on slides using Prolong Diamond Antifade Mountant (Thermo Fisher Scientific). The slides were imaged using a Zeiss LSM800 confocal microscope equipped with an Airyscan detector, using a 63x/1.4 oil objective in an Airyscan mode. Images were acquired using diode lasers 405, 561 and 670 nm in Z-stack mode. Image analysis was done using Fiji software version 2.1.0/1.53c.

The following primary antibodies were used in immunofluorescence stainings: Anti-Homer (rabbit, Proteintech # 12433-1-AP; 1:200; RRID: AB_2295573) Anti-Bassoon (mouse, SYSY #141011; 1:300; RRID: N/A) Anti-PSD95 (mouse, Thermo Fisher Scientific #MA1-045; 1:300; RRID: AB_325399) Anti-Map2 (rabbit, Proteintech # 17490-1-AP; 1:200; RRID: AB_2137880) Anti-NMDAR2 (rabbit, abcam # ab65783, 1:200; RRID: AB_1658870) Anti-GluA1 (mouse, SYSY #182-011, 1:200; RRID: AB_2113443) AntiGluN1 (mouse, SYSY #114-011, 1:500; RRID: AB_887750) Secondary antibodies (Thermo Fisher Scientific Scientific; all 1:1000): Goat anti-Mouse IgG (H+L) Cross-Adsorbed Secondary Antibody, Alexa Fluor™ 488 #A-11001 Goat anti-Rabbit IgG (H+L) Highly Cross-Adsorbed Secondary Antibody, Alexa Fluor™ 488 #A-11034 Goat anti-Rabbit IgG (H+L) Highly Cross-Adsorbed Secondary Antibody, Alexa Fluor™ 568 #A-11036 Goat anti-Mouse IgG (H+L) Highly Cross-Adsorbed Secondary Antibody, Alexa Fluor™ 568 #A-11031 Goat anti-Rabbit IgG (H+L) Cross-Adsorbed Secondary Antibody, Alexa Fluor™ 647 #A-21244 Goat anti-Mouse IgG (H+L) Cross-Adsorbed Secondary Antibody, Alexa Fluor™ 647 #A-21235

### Synapse analysis

Synapse analysis was performed based on immunofluorescence stainings, using the Fiji software, version 2.1.0/1.53c. Maximum intensity projections of Z-stacks were analyzed. Images were manually segmented to extract neuronal projections, and the number of Homer-Bassoon colocalizations and PSD-95 puncta were counted along 20 µm of neuronal projection, respectively. For the Gephyrin analysis, the puncta were counted from the cell body area. The data were visualized using the R/Bioconductor package *ggplot*.

### APOE ELISA

For the ELISA of iAstrocytes cultured alone, the cells were plated at a density of 15,000 cells per cm² in the wells of a 24-well plate. After seven days of routine culture, the media were collected. For co-cultures, media were collected at DIV14. All the collected media were spun to remove cell debris, and the quantity of APOE in the undiluted cell media was determined using an APOE ELISA kit (Thermo Fisher Scientific) according to the manufacturer’s instructions. Luminescence was measured using a Tecan M1000PRO device.

### Cholesterol measurement

To assess the total cholesterol concentration in cell media supernatants we used Cholesterol-GLO kit (Promega), according to instructions provided by the company. The supernatants were collected analogically to the APOE ELISA ones described above. Luminescence was measured using a Tecan M1000PRO device.

### APOE overexpression

APOE overexpression in iAstrocytes was achieved using lentiviral transduction. Lentiviral particles were prepared in HEK293T cells that had been transfected with either an APOE3-encoding plasmid (APOE_pLX307, Addgene #98317) or a “control” (“empty”) vector that had been obtained by removing the APOE3-encoding fragment using a restriction enzyme. The medium containing the lentiviral particles was collected 48 hours after transfection and concentrated using the Lenti-X Concentrator (Takara) according to the manufacturer’s instructions.

For lentiviral transduction, 400,000 iAstrocytes were plated in one well of a six-well plate. The concentrated lentiviral particles produced by the HEK293T cells in one well of the 6-well plate were added to the respective well of the iAstrocytes 24 hours after plating. Twenty-four hours after transduction, the iAstrocyte medium was fully exchanged for fresh medium. Forty-eight hours after transduction, puromycin selection was introduced. The puromycin-containing medium was fully exchanged daily up to 7 days post-transduction. The APOE level in the culture medium was then assessed using an ELISA, after which the cells were subjected to co-culture experiments.

### VT-107 (pan-TEAD) inhibition

For pan-TEAD inhibition, iAstrocytes were plated at the density of 5,000/cm^2^ on Matrigel-coated coverslips. 24 hours after plating, cells were treated with 3 μM V T-107 (MedChemExpress) in DMSO for 24 hours. Then, cells were collected and lysed for RNA isolation.

### RNA-seq

RNA collections from co-cultured cells were performed at DIV14. For RNA isolations cells were lysed directly in wells using TRI-Reagent (Merck). RNA was isolated with Direct-zol RNA MiniPrep kit (Zymo), according to manufacturer’s instructions. Sequencing libraries were prepared using KAPA mRNA HyperPrep kit (Roche). The library preparation procedure was performed according to the kit manufacturer’s instructions. For indexing, UMI in xGen™ UDI-UMI Adapters (IDT) were used. Library concentrations were determined using Quantus system (Promega) and library sizes were examined with TapeStation DNA ScreenTape & Reagents (Agilent) on TapeStation4200 Device (Agilent). Libraries were sequenced (2x100 bp, paired-end) using NovaSeq6000 device (Illumina).

### Computational analyses

#### Co-cultures RNAseq data processing

Raw RNA-seq reads were trimmed with TrimGalore ver. 0.6.7, using parameters ‘--paired -q 30 --stringency 3’. *BBsplit* was used to separate raw reads based on mapping against the consensus genome and the RatNor7 reference genome. Reads uniquely mapping to a single reference were retained, while ambiguously mapping reads were handled according to the analysis settings, enabling separation of reads for downstream analyses. Alignment against the consensus genome was implemented using STAR version 2.7.10 with default parameters and ‘outFilterMultimapNmax 1’. We used featureCounts version 2.0.3 with parameters ‘-p -O --countReadPairs -t exon -g gene_id’ to obtain per gene RNA-seq read counts. Annotation “Homo_sapiens.GRCh38.101.gtf” from Ensembl’s release version 101 was as a reference for samples aligned consensus genome and “Rattus_norvegicus.mRatBN7.2.107.gtf” for samples aligned for Rat genome.

### RNA sequencing data processing

RNA sequencing data from induced astrocytes (iAstrocytes) and rat hippocampal neurons maintained in co-culture were analyzed in R (v4.4.2) using Bioconductor packages. Gene annotations were retrieved from Ensembl using the biomaRt package by querying the *rnorvegicus_gene_ensembl* dataset (mRatBN7.2.113) for rat neuronal transcriptomes and the *hsapiens_gene_ensembl* (Hg38) dataset for human astrocyte transcriptomes. Duplicate entries were removed, and Ensembl gene identifiers were mapped to gene symbols, Entrez identifiers, and gene biotypes before downstream analyses.

### Differential gene expression analysis

Gene-level count matrices were imported into R, and differential gene expression was performed using R/Bioconductor package *DESeq2* (1.46.0). Raw counts were modeled using a negative binomial generalized linear model with size factor normalization and dispersion estimation.

To determine transcriptional differences between neurons exposed to astrocytes from different primate species, neurons co-cultured with human iAstrocytes were compared with neurons co-cultured with chimpanzee iAstrocytes, while accounting for experimental batch effects:

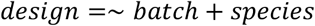

Genes were considered differentially expressed when they satisfied two thresholds: adjusted *P*-value < 0.1 and |log₂ fold change| > 0.59.

Normalized counts were obtained using the *counts* function in DESeq2, and variance-stabilized expression matrices were generated using the variance-stabilizing transformation (VST) for visualization and downstream analyses.

### Differential exon usage analysis

We computed differential exon usage (DEU) using R/Bioconductor package *DEXSeq* (1.54.1). RNA-seq reads were aligned to the reference genome (ratnor7, Rattus_norvegicus.mRatBN7.2.107.gtf) using STAR and the sorted BAM files were considered for further analysis. Gene annotations from the corresponding GTF file were used to construct non-overlapping exon-counting bins with the DEXSeq Python script, “*dexseq_prepare_annotation.py*”. Exon-level read counts compatible with *DEXSeq* were obtained using “*dexseq_count.py”*. The resulting count tables were imported into *R* and analyzed with the *DEXSeq* package to determine differentia exon usage (DEU) between human and chimpanzee samples using standard model settings and batch correction. We defined significant differential exon usage events based on adjusted *p*-values (padj < 0.05) and |LFC| >1.

### Analysis of astrocyte transcriptomes – identification of evolutionarily altered astrocyte genes (EAGs)

To identify species-specific transcriptional programs in astrocytes in co-cultures, RNA-seq profiles of human and chimpanzee iAstrocytes cultured with neurons were compared using *DESeq2* with the model:

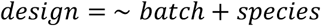

Genes significantly enriched in either species were identified using the thresholds described above. Differential expression results were visualized using volcano plots showing log_2_ fold changes versus -log_10_(*P*-values). Gene identifiers were converted to Entrez identifiers and gene symbols using Ensembl annotation tables retrieved via R/Bioconductor package *biomaRt*.

To evaluate evolutionary conservation of astrocyte transcriptional programs, iAstrocyte transcriptomes were compared with published fetal brain astrocyte datasets from human and macaque samples. This allowed us to identify evolutionarily affected genes (EAGs).

We assembled a pseudobulk transcriptome matrix of fetal astrocytes from published single-cell RNA sequencing datasets of human and macaque developing brains. To enable direct cross-species comparisons, single-cell reads were aligned to a consensus primate genome, following the strategy described in Ciuba et al., 2025, ensuring consistent gene models between human and macaque datasets.

Single-cell RNA sequencing datasets from fetal primate brains were analyzed using Seurat (v5). Cell type identities were assigned by reference mapping using the Azimuth framework with the fetal brain reference dataset (*fetusref*).

For each dataset, raw counts were converted into Seurat objects and normalized using *SCTransform*. Principal component analysis was performed using the top variable features. Shared features between query datasets and the reference atlas were identified, and cell identities were transferred using *FindTransferAnchors* followed by *MapQuery*.

Cell type annotations were assigned at two hierarchical levels using the reference annotations (*annotation.l1* and *annotation.l2*). Cells assigned to the level 1 astrocyte class were further filtered using marker-based module scores to enrich for bona fide astrocytes and reduce contamination from other major brain cell types. Module scores were calculated with *AddModuleScore* using astrocyte, neuronal, oligodendrocyte, OPC, vascular, microglial and radial glial marker sets. Astrocyte pseudobulk profiles were generated from cells annotated as astrocytes that ranked above the 80th percentile for the astrocyte module score and below the indicated exclusion thresholds for contaminating lineage scores: neuronal score below the 50th percentile, oligodendrocyte score below the 99th percentile and vascular score below the 99th percentile, with thresholds calculated across the annotated brain-cell subset. Raw RNA counts from retained astrocytes were summed gene-wise to generate one pseudobulk astrocyte profile per library.

Next, to obtain sample-level astrocyte expression profiles, we generated pseudobulk transcriptomes by summing raw counts across all cells annotated as astrocytes in the step above, within each library. Then, for each individual, the count matrix of the RNA assay was extracted, and counts corresponding to astrocyte-annotated cells were aggregated using row-wise summation. This procedure generated one pseudobulk astrocyte profile per library. The final dataset consisted of 16 human fetal astrocyte pseudobulk samples and 12 macaque fetal astrocyte pseudobulk samples.

Differential expression between human and macaque astrocytes was performed using R/Bioconductor package *DESeq2* (1.46.0) on the pseudobulk count matrix using the model:

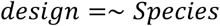

Genes were considered significantly differentially expressed when they satisfied the following criteria: adjusted p-value < 0.01 and |log_2_ fold change| > 0.59.

Genes upregulated in human astrocytes relative to macaque astrocytes were classified as human-enriched, whereas genes enriched in macaque astrocytes were classified as non-human primate–enriched. EAGs upregulated in human astrocytes were defined as genes upregulated in both comparison of human and chimpanzee iAstrocytes and human and macaque fetal astrocytes. EAGs downregulated in human astrocytes were defined as genes downregulated in both comparisons of human and chimpanzee iAstrocytes and human and macaque fetal astrocytes.

### Gene ontology enrichment

Functional enrichment analyses were performed using R/Bioconductor package *clusterProfiler*. Over-representation analyses were conducted separately for gene sets upregulated in neurons exposed to human astrocytes and those upregulated in neurons exposed to chimpanzee astrocytes.

Gene ontology enrichment was evaluated across the three Gene Ontology domains: Biological Process (BP), Cellular Component (CC), and Molecular Function (MF).

Enrichment significance was determined using Benjamini–Hochberg adjusted *P*-values (adjusted *P* < 0.01). Enrichment results were visualized using dot plots and gene networks generated using the R/Bioconductor package *enrichplot* package.

### Disease ontology enrichment

To assess potential associations with neurological disorders, rat genes differentially regulated in neurons exposed to primate astrocytes were mapped to human orthologs using *biomaRt*. Disease ontology enrichment was performed using the R/Bioconductor package *EnrichDO* package with human Entrez identifiers. The significance of enrichment was evaluated using adjusted *P*-values, and results were visualized using R/Bioconductor package *ggplot2*.

### Gene module and heatmap analyses

Selected differentially expressed genes related to extracellular matrix organization, mechanotransduction, lipid metabolism, ion homeostasis, and synaptogenic signaling were visualized using ComplexHeatmap together with circlize for color scaling.

Expression matrices were row-scaled (Z-score normalization) to highlight relative expression differences across species and experimental conditions. Gene sets were grouped into functional modules representing distinct astrocyte transcriptional programs and neuronal developmental pathways.

### Pan-TEAD inhibition RNA sequencing analysis

To examine transcriptional responses to inhibition of TEAD transcription factors, human iAstrocytes were treated with the pan-TEAD inhibitor VT107 for 24 hours. RNA-seq count matrices were analyzed using *DESeq2* (1.46.0) with the model:

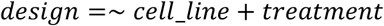

Differential expression was determined using the Wald test with Benjamini–Hochberg correction. Genes with adjusted p-values < 0.1 and |log₂FC| > 0.59 were considered significant.

Gene ontology enrichment of TEAD-responsive genes was performed using R/Bioconductor package *clusterProfiler*, and enriched functional categories were visualized using dot plots and enrichment networks.

### Analysis of human-chimpanzee sequence divergence of APOE promoter and enhancer

All sequence handling and motif analyses were performed in *R* using R/Bioconductor packages *Biostrings*, *TFBSTools*, *JASPAR2022*, *dplyr*, and *ggplot2*. To assess the impact of evolutionary sequence divergence on predicted transcription factor binding in the APOE enhancer, we analyzed orthologous 1 kb human and chimpanzee sequences corresponding to the regulatory element interval of interest (promoter or the looped enhancer, we considered the region centered on the ATAC-seq peak displaying the highest activity in MPRA that is the chimpanzee peaks). Pairwise global alignments between human and chimpanzee sequences were generated using the *pairwiseAlignment* function from the R/Bioconductor package *Biostrings* package. Alignment tables were then constructed to map nucleotide positions between species while preserving indels. Position frequency matrices from the JASPAR2022 CORE vertebrate collection were retrieved using *getMatrixSet* and converted to position weight matrices (PWMs) using *TFBSTools::toPWM*.

Motif scanning was performed separately in two reciprocal modes:

(1) In the human-centered analysis, candidate TF binding sites were identified in the human sequence using R/Bioconductor package *Biostrings::matchPWM* on both forward and reverse strands. For each human motif hit, the orthologous chimpanzee sequence was extracted from the human-chimpanzee alignment.

(2) In the chimpanzee-centred analysis, the same procedure was applied in reverse: candidate sites were first identified in the chimpanzee sequence and the orthologous human sequence was then extracted from the alignment. This reciprocal design was used to reduce directional bias introduced by motif discovery in only one species.

For each matched human-chimpanzee site, raw PWM scores were recalculated directly on the exactly aligned motif sequences using R/Bioconductor package *Biostrings*::*PWMscoreStartingAt*. Reverse-strand sites were reverse-complemented prior to rescoring so that both species were evaluated in the correct motif orientation. To facilitate comparison across motifs with different score ranges, raw scores were normalized to the motif-specific PWM range using the formula:

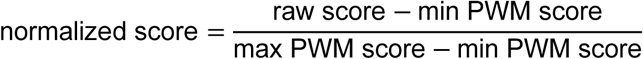

where the minimum and maximum PWM scores were computed as the sums of the minimum and maximum column-wise PWM values, respectively.

For the human-centered analysis, species differences were quantified as:

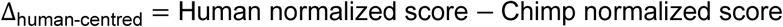

For the chimpanzee-centered analysis, species differences were quantified as:

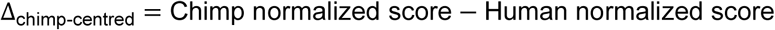

Sites overlapping alignment gaps in the orthologous sequence were classified as indel-disrupted and excluded from quantitative cross-species score comparison. We retained only sites that satisfied all of the following criteria: no indel disruption, at least one nucleotide difference between the human and chimpanzee motif-aligned sequences, a normalized motif score greater than 0.84 in at least one species, an absolute normalized score difference greater than 0.05, and motif assignment to a transcription factor expressed in astrocytes. To reduce redundancy in summary plots, only the strongestfor eachte per motif was retained separately in the human-stronger and chimpanzee-stronger sets.

### Calcium imaging quantification and statistical analysis

Single-cell calcium traces from neurons co-cultured with human iAstrocytes transduced with empty vector or APOE were analyzed in R using a custom pipeline. After excluding the first 100 frames, (ΔF/F) was calculated for each cell using a rolling baseline defined as the 10th percentile within a 100-frame sliding window. Calcium peaks were detected as local maxima above the 80th percentile of the trace, requiring a minimum signal drop of 0.10 (ΔF/F) units on both sides within a 15-frame window. Peaks occurring within 20 frames were merged by retaining the higher event. The total number of peaks per cell was used as a measure of neuronal activity. To test the effect of APOE overexpression, peak counts were analyzed using a negative binomial mixed-effects model with treatment as a fixed effect and iPSC line and experiment as random intercepts. Effect sizes were expressed as fold change relative to empty-vector controls by exponentiation of the model coefficient.

### Massively Parallel Reporter assay (MPRA)

MPRA was performed according to a modified protocol by Tewhey et al, (2016),^135^ described in detail below. The library was ordered from Twist Biosciences as a lyophilized oligonucleotide pool. Each oligonucleotide was designed as a 200-basepair stretch of DNA flanked by universal amplification adapters. Final library structure followed the pattern: ACTGGCCGCTTGACG-[tested 200 bp enhancer sequence]-CACTGCGGCTCCTGC. Upon receipt, DNA was spun down and suspended in UltraPure™ DNase/RNase-Free Distilled Water (Invitrogen) to a final concentration of 10ng/μl. The pool was diluted to 1ng/μl. The resuspended library was stored at −80°C.

Library amplification was carried out in emulsion PCR. For a library of approximately 6000 fragments, 3 separate emulsion PCR reactions were prepared on ice, each comprising: 0.5μM of primer R20_MPRA_v3_F, 0.5μM of primer R19_MPRA_v3_20I_R 0.5μM, 1.86ng of oligonucleotide library, 25μl of New England BiolabsNext Ultra II Q5® Master Mix (New England Biolabs), 0.5μl of Q5® High-Fidelity DNA Polymerase (New England Biolabs), 2ng of BSA, UltraPure™ DNase/RNase-Free Distilled Water was used to bring the reaction volume to 50μl. For each 50μl of reaction, the following reagents were added to create an emulsion: 220μl of TegoSoft DEC (Evonik), 60μl of ABIL WE (Evonik), 20μl of Mineral Oil (SIGMA), for a total of 300μl per single PCR reaction mix. To emulsify the reaction, the mixture was then vortexed for 5’ at 4°C. The mixture was then distributed into 96 well plate on ice, in 50μl portions. The plate was then incubated ia n thermocycler according to the following conditions: step 1: 95°C, 30 seconds, step 2: 95°C, 20 seconds, step 3: 60’°C, 10 seconds, 72°C, step 4: 15 seconds, return to step 2 35 times, step 5: 72°C, 5 minutes. The emulsion was subsequently broken up by the addition of 1ml of 2-butanol (Thermo Fisher Scientific), 50μl of Agencourt AMPure XP beads (Beckman Coulter), 80μl of binding buffer (2.5M NaCl, 20% PEG-8000 (Promega) per 350μl of emulsion mix, followed by vigorous vortexing. The mixture was then incubated at room temperature for 10 minutes and spun for 5 minutes at 2,900xg. The resulting solution contained two clearly visible phases. The upper, organic phase was carefully collected and discarded. The remaining solution was then placed on a magnetic rack for 20 minutes. Supernatant was removed, and beads were washed once with 1 ml of 2-butanol, and three times with freshly made 80% ethanol. The beads were subsequently dried at room temperature for 5 minutes and eluted using elution buffer (EB, Qiagen). The resulting concentration of DNA was estimated using NanoDrop. Size distribution was tested using TapeStation. The barcoded library was cloned into the destination plasmid pMPRAv3:Δluc:ΔxbaI linearised by digestion with SfiI restriction enzyme using GeneArt™ Gibson Assembly HiFi Master Mix (Invitrogen). Ligation was carried out by incubating the mixture for 60 minutes at 50°C, followed by SPRI purification using Agencourt AMPure XP beads (Beckman Coulter), according to the manufacturer’s instructions using a 1.8:1 bead:sample ratio. Ligated plasmid was eluted from beads in 20μl of elution buffer (EB, Qiagen) and quantified using NanoDrop. 25ng of the solution was then transformed per 50μl of 10-beta Electrocompetent E. Coli (New England Biolabs) by electroporation (Gene Pulser/MicroPulser Electroporation Cuvettes, 0,2cm gap, (BioRad), electroporator: electroporation settings 2kV, 200ohm, 25uF). Serial dilution was performed after electroporation to estimate the number of CFU.

After cloning the first sequencing step to determine the association between oligo sequence and barcode was performed. Library purification consisted of two rounds of PCR, each followed by SPRI purification.

For the first step of library preparation, two 50μl PCR reactions were set up on ice as follows: 200ng of plasmid library prepared in the previous step, 50μl of New England BiolabsNext Ultra II Q5® Master Mix (New England Biolabs), 0,5 μM of R45_MPRA_v3_Amp2Sa_Illu_FR primer, 0,5 μM of R46_Illu_univ_adapter; UltraPure™ DNase/RNase-Free Distilled Water was used to bring the reaction volume to 100μl. The reaction mixture was then incubated in a thermocycler according to the following conditions: step 1: 95°C, 20 seconds. step 2: 95°C, 20 seconds, step 3: 62°C, 15’ seconds, step 4: 72°C for 30 seconds, return to step 2, 8x, step 5: 72°C, 2 minutes. The mixture was then purified using AMpure XP beads according to the manufacturer’s protocol for purification of small fragments, using 0.6:1 bead:sample ratio. DNA was eluted with 15μl of EB. Size distribution was examined with TapeStation with the product peak expected at 365bp.

For the second step of library preparation, index-containing primers were used (primers S69-S70, T17-T23, T36, Z61-Z67, Z70-Z71). The PCR mixture was set-up on ice as follows: 10μl of eluted DNA from the first amplification step, 50μl of New England BiolabsNext Ultra II Q5® Master Mix (M0544S), 0,5μM of barcoding primer (one of primers S69-S70, T17-T23, T36, Z61-Z67, Z70-Z71), 0,5μM of R46_Illu_univ_adapter, UltraPure™ DNase/RNase-Free Distilled Water was used to bring the reaction to 100μl. The reaction mixture was then incubated in a thermocycler according to the following conditions: step 1: 95°C, 20 seconds. step 2: 95°C, 20 seconds, step 3: 64°C, 30 seconds, step 4: 72°C for 30 seconds, return to step 2, 6x, step 5: 72°C, 2 minutes. The mixture was then purified using Agencourt AMPure XP beads according to manufacturer’s protocol, using 1.8:1 bead:sample ratio. DNA was eluted with 15μl of elution buffer (EB, Qiagen). Size distribution was examined with TapeStation, with the product peak expected at 397bp. Libraries were sequenced (2x150 bp, paired-end) using NovaSeq6000 device (Illumina).

Subsequently, the minimal promoter-GFP cassette was cloned into the library. The insert cassette was amplified from plasmid Addgene 109036 as follows: 0,5μM of primer R21_ED_MPRA_GFP_F, 0,5μM of primer R22_ED_MPRA_GFP_R, 0,2μM of dNTPs (ThermoFisher Scientific). 5x Q5 buffer (New England Biolabs), 10μl, Q5 High-Fidelity DNA Polymerase (New England Biolabs), 0,5μl, plasmid 50ng, UltraPure DNase/RNase-Free Distilled Water was used to bring the reaction to 50μl. The reaction mixture was then incubated in a thermocycler according to the following conditions: step 1: 98°C, 30 seconds. step 2: 98°C, 10 seconds, step 3: 69°C, 30 seconds, step 4: 72°C for 30 seconds, return to step 2, 35x, step 5: 72°C, 2 minutes, and purified using AMPure XP beads according to manufacturer’s protocol for purification of small fragments, using 0.65:1 bead:sample ratio. DNA was eluted in elution buffer (Qiagen).

The library was linearised using AsiSI (New England Biolabs). Minimal promoter-GFP cassette was cloned into the library using 2x GeneArt™ Gibson Assembly HiFi Master Mix (Invitrogen), incubated for 90 minutes at 50°C. The mixture was then purified using Agencourt AMPure XP beads (Beckman Coulter) according to manufacturer’s protocol, using 1.8:1 bead:sample ratio. DNA was eluted with 25μl of buffer EB (Qiagen). In order to remove uncut plasmid, purified DNA was treated AsiSI (New England Biolabs) and Exonuclease V and purified using Agencourt AMPure XP beads (Beckman Coulter) according to manufacturer’s protocol, using 1.8:1 bead:sample ratio, eluted in 20μl of EB (Qiagen) and quantified using NanoDrop.

The plasmid library was then transformed into 10-beta Electrocompetent E. Coli (New England Biolabs) by electroporation (Gene Pulser/MicroPulser Electroporation Cuvettes, 0,2cm gap, (BioRad), electroporator: electroporation settings 2kV, 200ohm, 25uF). Serial dilution was performed after electroporation to estimate the number of CFU.

### Cell transfection, cDNA, and MPRA readout library preparation

Cells were seeded at a density of 75000 cells/cm2 of petri dish the day before transfection, in 75cm2 cell culture bottles. Line LN229 was seeded directly on cell-culture-treated plastic dishes. On the day of transfection, the medium was changed approximately 30 minutes before transfection; 7.8ml of fresh growth medium was added. For each 75cm^2^ bottle, transfection was performed as follows: in two separate tubes, DNA mix (1972.5μl of Opti-MEM Reduced-Serum Medium, (ThermoFisher Scientific), 47.5 ug of plasmid library) and lipofectamine mix (1972.5μl of Opti-MEM Reduced-Serum Medium, 157.5μl of Lipofectamine Stem Transfection Reagent). DNA mixture was then added to the Lipofectamine mixture, mixed gently, and incubated at room temperature for 10 minutes before being added to the cells drop by drop. One day after transfection, the medium was changed. Cells were collected on the second day after transfection. First, the medium was removed and kept. Then, cells were washed once with PBS, and the wash-out was kept. Then, Accutase (SIGMA) was added. Spent medium wash then spun for 5 minutes at 500g, and discarded, leaving the pellet. Then, the PBS wash-out was added to the same tube, and spun as before, and supernatant was discarded. After cells were detached, Accutase was collected, added to the pellet of previously collected cells, and the plate was once more washed with PBS, and the washout was collected in the same tube. The tube was then spun for 5 minutes at 500g. Most of the supernatant was then collected and split into two equal portions in 1.5 Eppendorf tubes. Each portion was spun (as above) and washed two more times with 1ml of PBS. After the final spin, as much of the supernatant as possible was removed, and cell pellets were frozen.

The cDNA synthesis was performed using the SuperScript III Reverse transcriptase kit (Life Technologies). First, RNA was extracted from each of the aliquots of frozen cells using Direct-zol™ RNA MiniPrep (Zymo) kit according to the manufacturer’s instructions. RNA was stored at −80°C. cDNA reaction was set-up on ice as follows: 2 pmol of primer R44_MPRA_v3_Amp2Sc_R, 500ng of RNA, 1μl of 10 mM dNTP Mix (ThermoFisher Scientific), UltraPure water (Invitrogen) to 10μl. The mixture was then incubated at 65°C for 5 minutes, and then on ice for at least 1 minute. Contents of the tube was collected by brief centrifugation. In ice, the following components were added to the reaction 4μl of 5X First-Strand Buffer (from Super Script II kit, Life Technologies), 1μl of 0.1 M DTT (from Super Script II kit, Life Technologies), 1μl of 1μl of SUPERase•In™ RNase Inhibitor, 20 U/μl (ThermoFisher Scientific), 1μl of SuperScript™ III RT (Life Technologies). The reaction was mixed gently by pipetting up and down, and incubated for 80 minutes at 47°C, and subsequently inactivated by incubation for 15minutes at 70°C. Purified using Agencourt AMPure XP beads (Beckman Coulter)) according to manufacturer’s protocol, using 1.8:1 bead:sample ratio, and eluted in 10μl of UltraPure water (Invitrogen).

To standardise library concentrations and minimise amplification bias, samples were amplified by qPCR to estimate the relative concentration of GFP cDNA, cDNA, or plasmid library diluted 500x was used as a template. The following reaction was set up on ice: 1μl of cDNA, diluted 2x or plasmid library, diluted 500x. 5μl of New England BiolabsNext Ultra II Q5^®^ Master Mix (New England Biolabs), 1μl of SYBR™ Green I Nucleic Acid Gel Stain (ThermoFisher Scientific, diluted 1:10000, R48_MPRA_v3_Illu_GFP_F to concentration of 0.5μM, R46_Illu_univ_adapter to concentration of 0.5μM (of 10μM), UltraPure water was used to bring the reaction to a total volume of 10μl. The reaction mix was protected from light in Hard-Shell® 384-Well PCR Plates (Bio-Rad). Plates were then vortexed and spun. qPCR reaction was carried out in Opus Real-Time PCR System according to the following conditions: step 1: 95°C, 20 seconds. step 2: 95°C, 20 seconds, step 3: 65°C, 20 seconds, step 4: 72°C for 30 seconds, return to step 2, 40x, step 5: 72°C, 2 minutes. cDNAs and plasmid solutions were diluted to march the relative concentration of the most dilute sample.

For the first step of library preparation, a 50μl PCR reaction was set up on ice for each sample, as follows: 2.5μl of cDNA/plasmid library diluted to match the concentration of the most dilute sample. 25μl of New England BiolabsNext® High-Fidelity 2X PCR Master Mix (New England Biolabs), primer R45_MPRA_v3_Amp2Sa_Illu_F to a final concentration of 0,5μM, primer R46_Illu_univ_adapter to a final concentration of 0,5μM, UltraPure water was used to brin the total reaction volume to 50μl. The reaction mixture was then incubated in a thermocycler according to the following conditions: step 1: 95°C, 20 seconds. step 2: 95°C, 20 seconds, step 3: 65°C, 20’ seconds, step 4: 72°C for 30 seconds, return to step 2, 10x(11cycles in total), step 5: 72°C, 2 minutes. The mixture was then purified using AMpure XP beads according to manufacturer’s protocol for purification of small fragments, using 0.6:1 bead:sample ratio. DNA was eluted with 15μl of buffer EB (Qiagen).

For the second step of library preparation, index-containing primers were used (primers S69-S70, T17-T23, T36, Z61-Z67, Z70-Z71, see primer list). The PCR mixture was set-up on ice as follows: 10μl of eluted DNA from the first amplification step, 50μl of New England BiolabsNext® High-Fidelity 2X PCR Master Mix (New England Biolabs), barcoding primer (one of primers S69-S70, T17-T23, T36, Z61-Z67, Z70-Z71) to concentration of 0,5μM of, primer R46_Illu_univ_adapter to concentration of 0,5μM, UltraPure water to 100μl. The reaction mixture was then incubated in a thermocycler according to the following conditions: step 1: 95°C, 20 seconds. step 2: 95°C, 20 seconds, step 3: 64°C, 30 seconds, step 4: 72°C for 30 seconds, return to step 2, 7x (in total, 8 cycles), step 5: 72°C, 2 minutes. The mixture was then purified using Agencourt AMPure XP beads (Beckman Coulter) according to manufacturer’s protocol, using 1.8:1 bead:sample ratio. DNA was eluted with 15μl of buffer EB (Qiagen). Size distribution was examined with TapeStation with the product peak expected at 268bp. Libraries were sequenced (2x150 bp, paired-end) using NovaSeq6000 device (Illumina).

### MPRA data processing

Analysis was conducted as described by Tewhey et. al, (2016). Paired-end 150 bp reads from the sequencing of the mpraΔorf library were merged into single amplicons using Flash v1.2.7 (flags: -r 150, -f 220, -s 10). Amplicon sequences were kept if the 5’ adapter matched with a Levenshtein distance of 3 or less, and 2 bp at the edges of both the 5’ and internal constant sequences matched perfectly. Oligo sequences from the passing reads were then mapped back to the expected oligo sequences using BWA mem version 0.7.9a (flags: -L 100 -k 8 -O 5). Alignment scores were calculated as matching bases divided by the expected oligo size and reads with alignment scores of less than 0.95 were discarded. Remaining oligo/barcode pairs were then merged and barcodes attributable to multiple oligo sequences were marked as conflicting and removed from further analysis. Barcodes matched with more than one enhancer were then removed using an R script, and barcodes were converted to reverse complements. The final association library consisted of a fasta list of enhancer candidates with individual barcodes.

Analysis was conducted similarly to the method described in Tewhey et. al, (2016). Only R1 reads were considered. Reads were trimmed to 30 bp using cutadapt (flags −l 30 -o <SAMPLE_NAME>.fastq). Subsequently, they were filtered according to the Levenshtein distance of 4 or less was required within the constant sequence at the end of the tag-seq read with the two bases directly adjacent to the barcode (base 21 & 22) required to match perfectly. The output was then trimmed again, this time removing 11bp at the 5’ end using cutadapt (flags −l 11 -o <SAMPLE_NAME>.fastq).

Sequences were then aligned to a FASTA list of barcodes generated previously. Alignment was performed using bwa (flags: -L 0 -k 19 -T 0). Relevant lines were then extracted using samtools (samtools view | cut -f 3 | grep _v “\*” >

<OUTPUT>.txt and samtools view | cut -f 3, 10 | grep -v “\*” >

<OUTPUT>.txt

For each library, the sum of all aligned barcodes associated with a given CRE was treated as counts, and analysed using the R/Bioconductor package *DESeq2*.

## Supplementary Figures

**Supplementary Figure 1.**
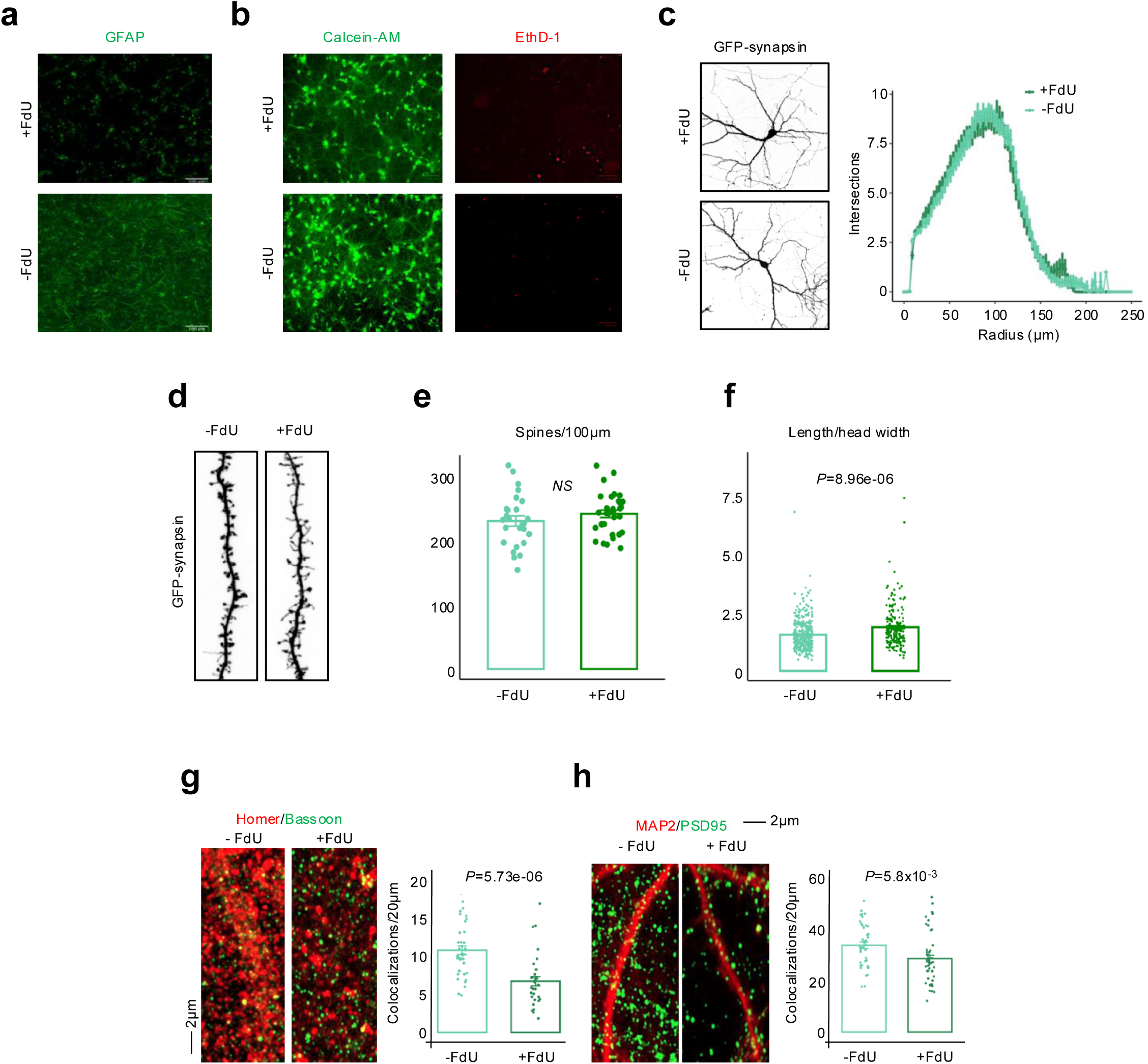
Effect of astrocyte depletion on primary rat hippocampal neurons. **a** Endogenous rat astrocytes were eliminated from dissociated hippocampal cultures by treatment with the antimitotic agent 5-fluoro-2′-deoxyuridine (FdU). Astrocyte depletion was confirmed by GFAP immunostaining in DIV14 cultures. Control cultures (-FdU) contain endogenous astrocytes. **b** FdU treatment does not markedly affect neuronal viability in D14 neuronal cultures. Live and dead cells were visualized using Calcein-AM (green) and Ethidium homodimer-1 (EthD-1; red) staining, respectively. **c** Representative morphology of DIV14 rat hippocampal neurons expressing GFP-synapsin cultured in the presence or absence of FdU. Sholl analysis comparing the neuronal morphology in the absence and absence of FdU did not show a significant difference between conditions (right panel). **d** Representative images of dendritic segments from GFP-synapsin-expressing neurons illustrating dendritic spine morphology in FdU-treated and control D14 neuronal cultures. **e** Quantification of dendritic spine density (spines per 100μm dendrite) shows no significant difference between conditions (*P* – Student’s t-test). **f** Analysis of dendritic spine morphology using the spine length-to-head width ratio reveals a shift toward shorter and thicker spines in cultures containing astrocytes (-FdU) compared with astrocyte-depleted conditions (+FdU) (*P* – Student’s t-test). **g** Left: Representative immunofluorescence images of Homer (postsynaptic; red) and Bassoon (presynaptic; green) puncta in neurons cultured in the presence or absence of astrocytes. Right: Quantification of Homer-Bassoon colocalizations along dendrites (per 20 μm) indicates reduced excitatory synapse maturation following astrocyte depletion. **h** Left: Representative images of PSD95 (green) puncta along MAP2-positive dendrites (red). Right: Quantification shows decreased density of PSD95-positive excitatory synapses in FdU-treated cultures. For panels f-g: N experiments = 3; n cells = 32 (+FdU) and 26 (-FdU); N spines = 7,650 (+FdU) and 6,484 (-FdU). For panels H-J, each dot represents synaptic puncta quantified along 20μm of dendritic length (N experiments = 3). Scale bars: 2 μm (E, H–J).

**Supplementary Figure 2.**
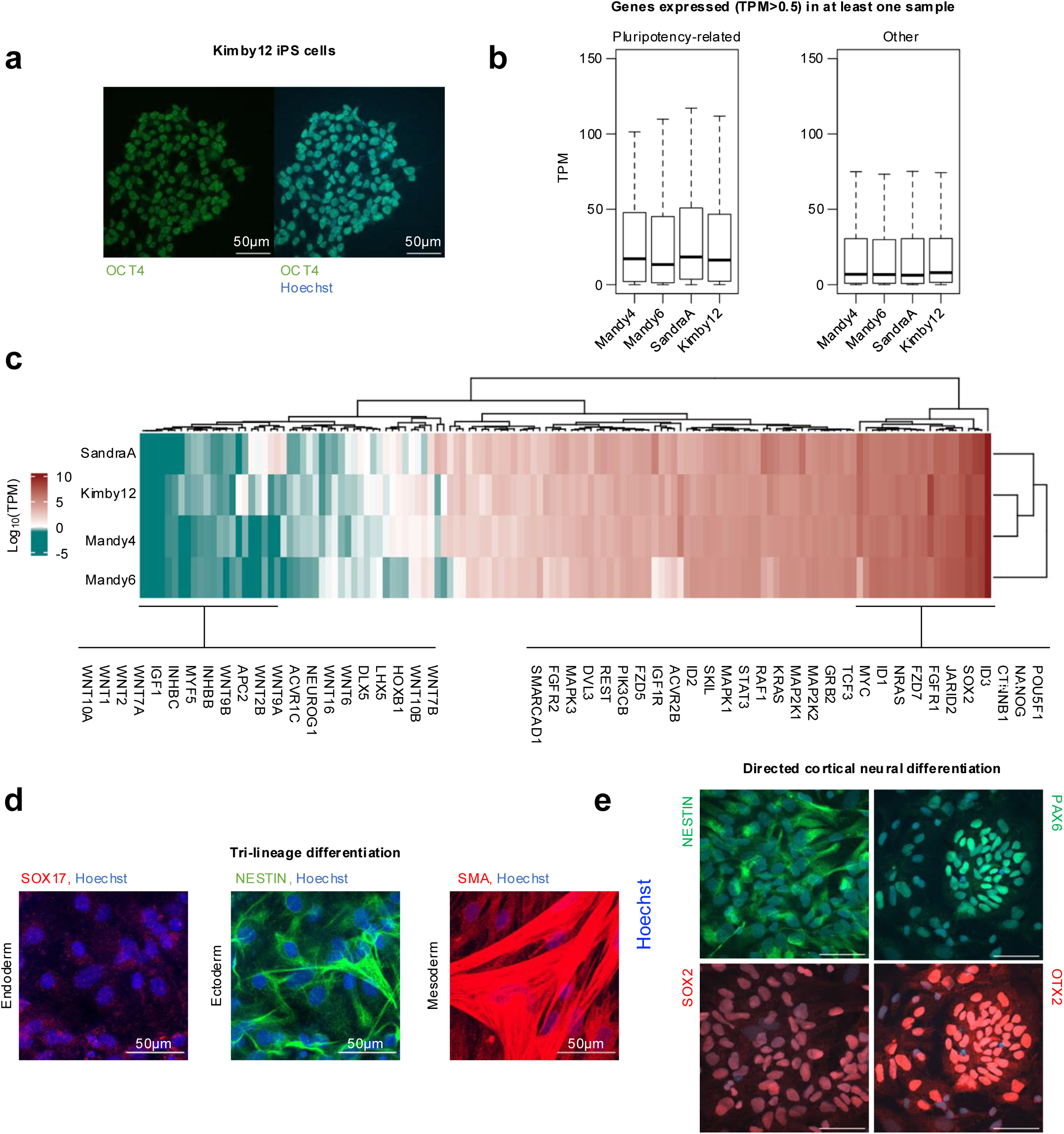
Validation of Kimby 12 induced pluripotent stem (iPS) cells. **a** Kimby12 iPS cells express Oct4, the canonical marker of pluripotent stem cells. Representative image of an iPS cell colony assayed with immunofluorescence for OCT4 expression (green). DNA was counterstained with Hoechst. **b** Expression levels of pluripotency-associated and non-pluripotency genes across iPS cell lines, including Kimby12, assessed by RNA-seq (TPM > 0.5 in at least one sample), confirming maintenance of a pluripotent transcriptional program comparable to established lines. **c** Hierarchical clustering of expression (log_2_ TPM) of pluripotency and iPS-cell differentiation-related genes across iPS cell lines, showing enrichment of pluripotency-associated genes and low expression of differentiation-associated genes in Kimby12 cells. **d** Spontaneous differentiation of Kimby12 iPS cells to three germ layers through the generation of embryoid bodies. Kimby 12 iPS cells were differentiated into ectoderm, endoderm, and mesoderm, as confirmed by immunofluorescence staining for the respective markers. **e** Patterned differentiation of Kimby12 iPS cells to cortical neural progenitors (iNPs). The Kimby12 iNPs express PAX6, OTX2, Nestin, and SOX2 proteins, the canonical markers of neural progenitor identity in primates. All scale bars: 50μm.

**Supplementary Figure 3.**
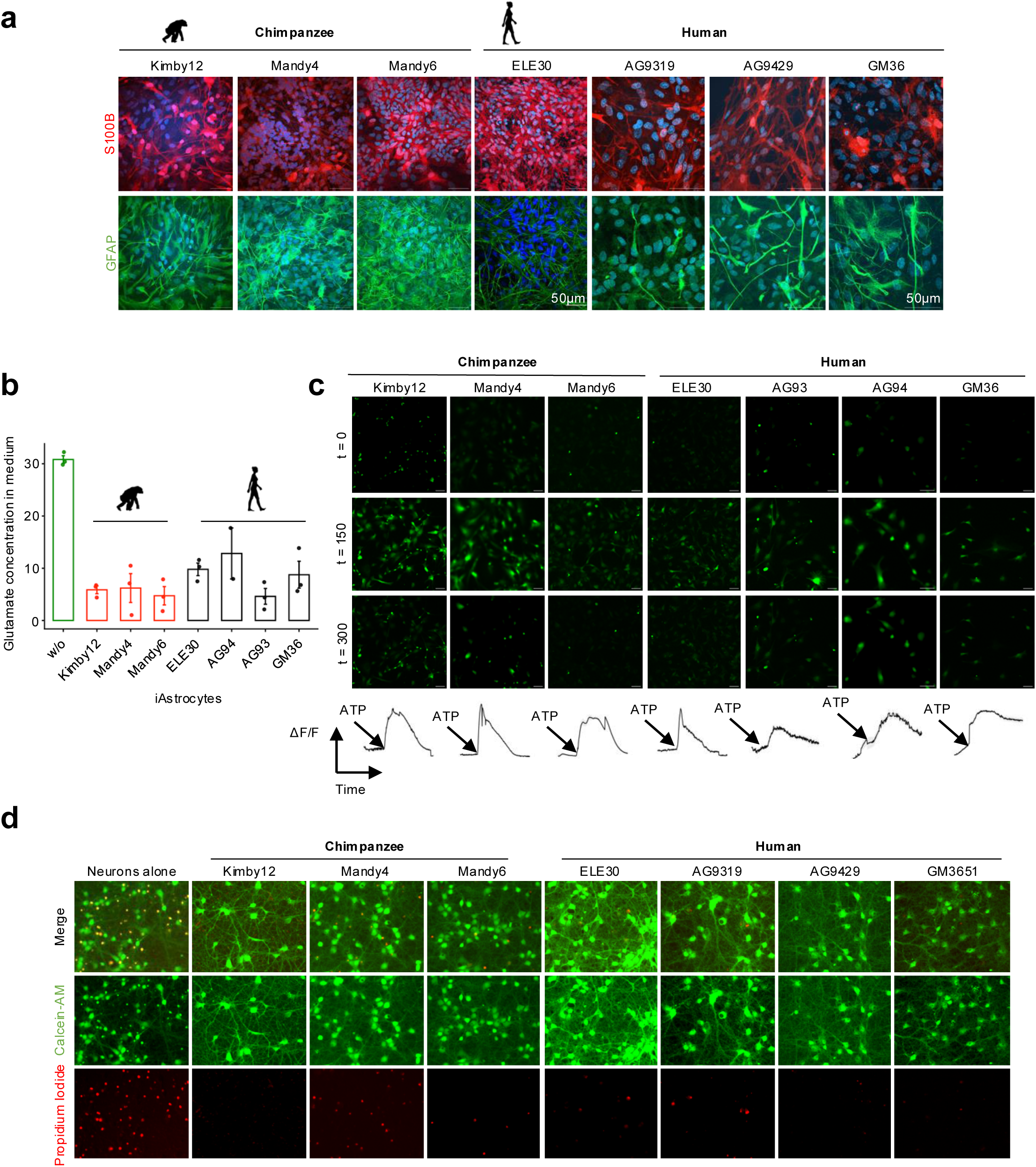
Validation of primate iAstrocytes used in the study. **a** Representative immunofluorescence images of chimpanzee and human induced astrocytes (iAstrocytes) expressing canonical astrocyte markers S100B (red) and GFAP (green). Nuclei are stained with DAPI (blue). Scale bar: 50μm. **b** Uptake of extracellular glutamate by primate iAstrocytes. Glutamate concentration in the culture medium was measured after incubation with chimpanzee or human iAstrocyte lines. “w/o” indicates medium without cells. Data represent mean ± SEM from n=3 independent experiments. **c** ATP-induced calcium responses in primate iAstrocytes. Representative fluorescence images show intracellular calcium signals at t = 0, 150, and 300 s following ATP stimulation. Traces below depict average ΔF/F calcium responses over time for each iAstrocyte line (n=50 cells per line). **d** Primate iAstrocytes support survival of primary rat hippocampal neurons. Representative images of neurons cultured alone or in co-culture with chimpanzee or human iAstrocytes. Calcein-AM (green) labels live neurons, whereas propidium iodide (red) marks dead cells.

**Supplementary Figure 4.**
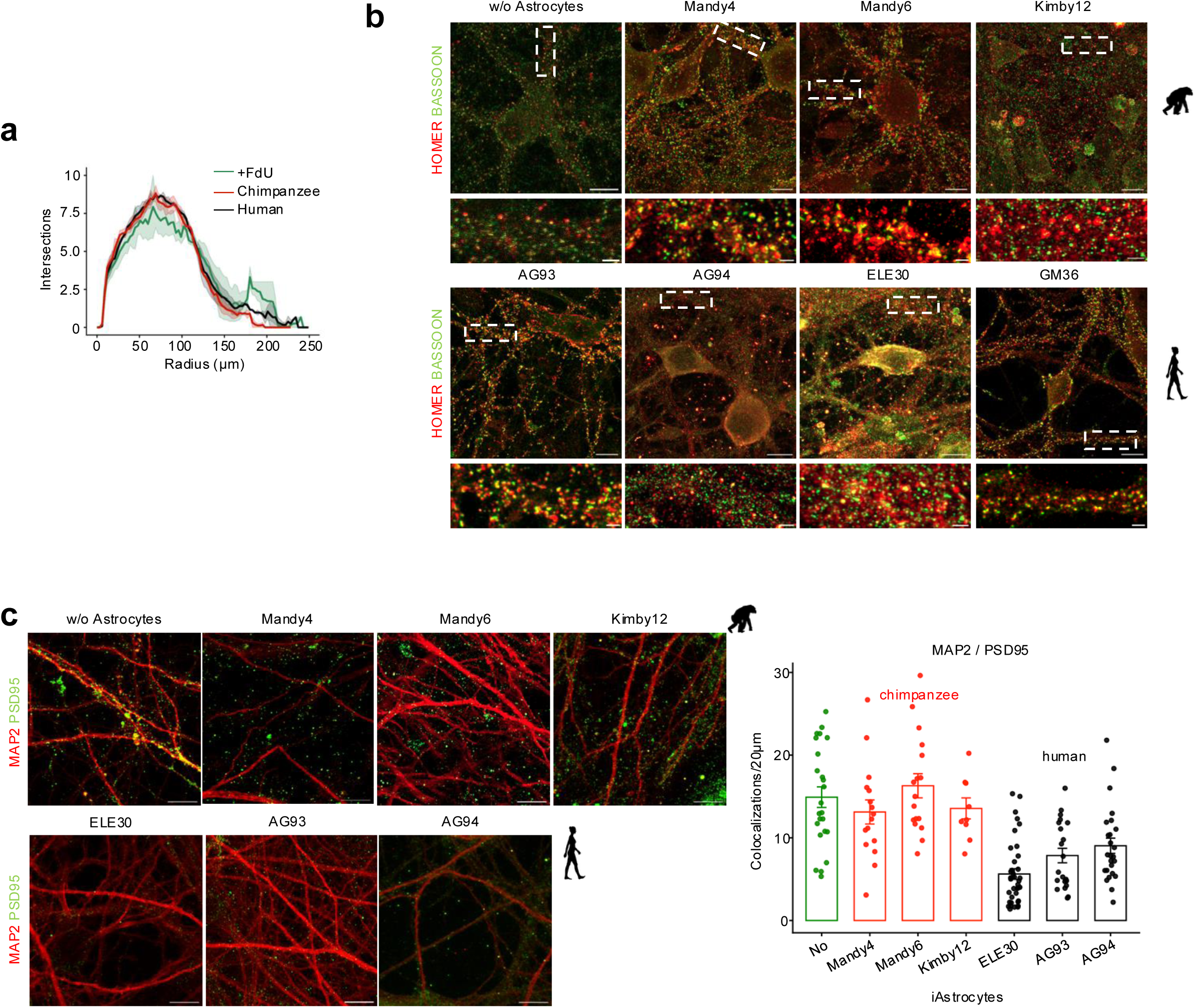
Human astrocytes alter excitatory synapse maturation without changing synapse number or dendritic architecture. **a** Sholl analysis of dendritic complexity in rat hippocampal neurons cultured alone or in co-culture with chimpanzee or human iAstrocytes. No significant differences in dendritic branching were observed across conditions; n cells = 24 (control – rat neurons cultured in the presence of FdU), 49 (HS; ELE30=20, AG94=29), 44 (PT; Mandy4=26, Mandy6=18) **b** Representative immunofluorescence images of Homer (postsynaptic; red) and Bassoon (presynaptic; green) puncta in rat hippocampal neurons cultured either alone or in the presence of chimpanzee or human iAstrocytes. Insets show magnified regions (dashed boxes). N experiments = 3. Scale bars: 50μm (overview), 2μm (insets). **c** Representative images corresponding to Figure 2D showing PSD95 (green) puncta along MAP2-positive dendrites (red) in rat neurons cultured either alone or with primate iAstrocytes. N experiments = 3. Scale bars: 50μm (overview), 2μm (insets), and quantification of PSD95 puncta along MAP2 dendrites in individual cell lines. Each dot represents one projection measurements (n experiments = 3).

**Supplementary Figure 5.**
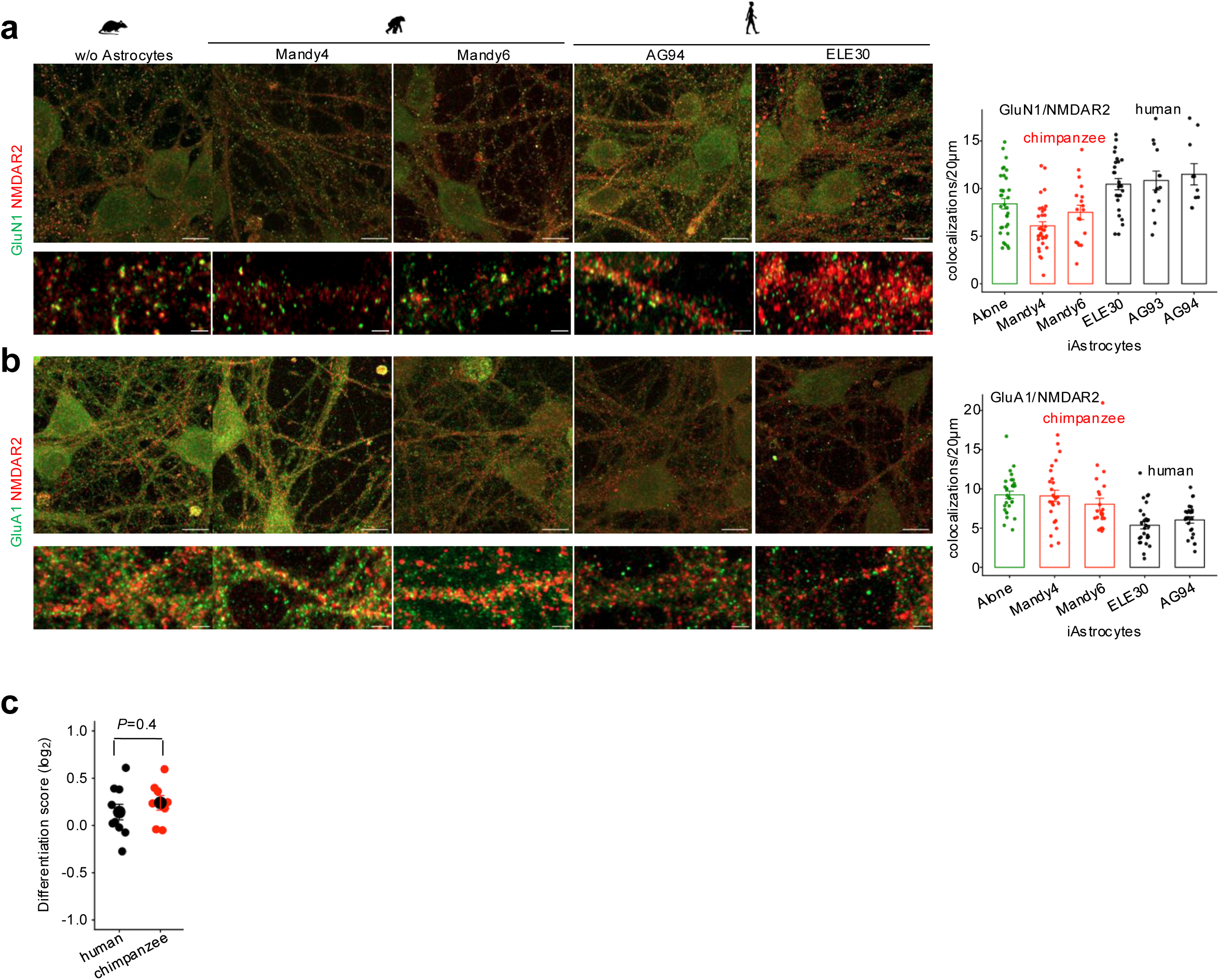
Additional synaptic quantifications in neurons cultured with primate iAstrocytes. **a** Right: Representative immunofluorescence images of NMDA receptor-containing synapses (GluN1, green; NMDAR2, red) in neurons cultured alone or with primate iAstrocytes. Left: Quantification of GluN1-NMDAR2 colocalizations in neurons cultured alone or with primate iAstrocytes (as in Figure 2F), presented by individual iAstrocyte cell line. Each dot represents synaptic puncta quantified along 20μm of dendritic length. **b** Right: Representative immunofluorescence images of NMDA receptor-containing synapses (GluA1, green; NMDAR2, red) in neurons cultured alone or with primate iAstrocytes. Left: Quantification of GluA1-NMDAR2 colocalizations in neurons cultured alone or with primate iAstrocytes (as in Figure 2F), presented by individual iAstrocyte cell line. Each dot represents synaptic puncta quantified along 20μm of dendritic length. **c** Differentiation score of iAstrocytes in co-culture with rat neur The differentiation score was computed as in Ciuba *et al.* (2025).

**Supplementary Figure 6.**
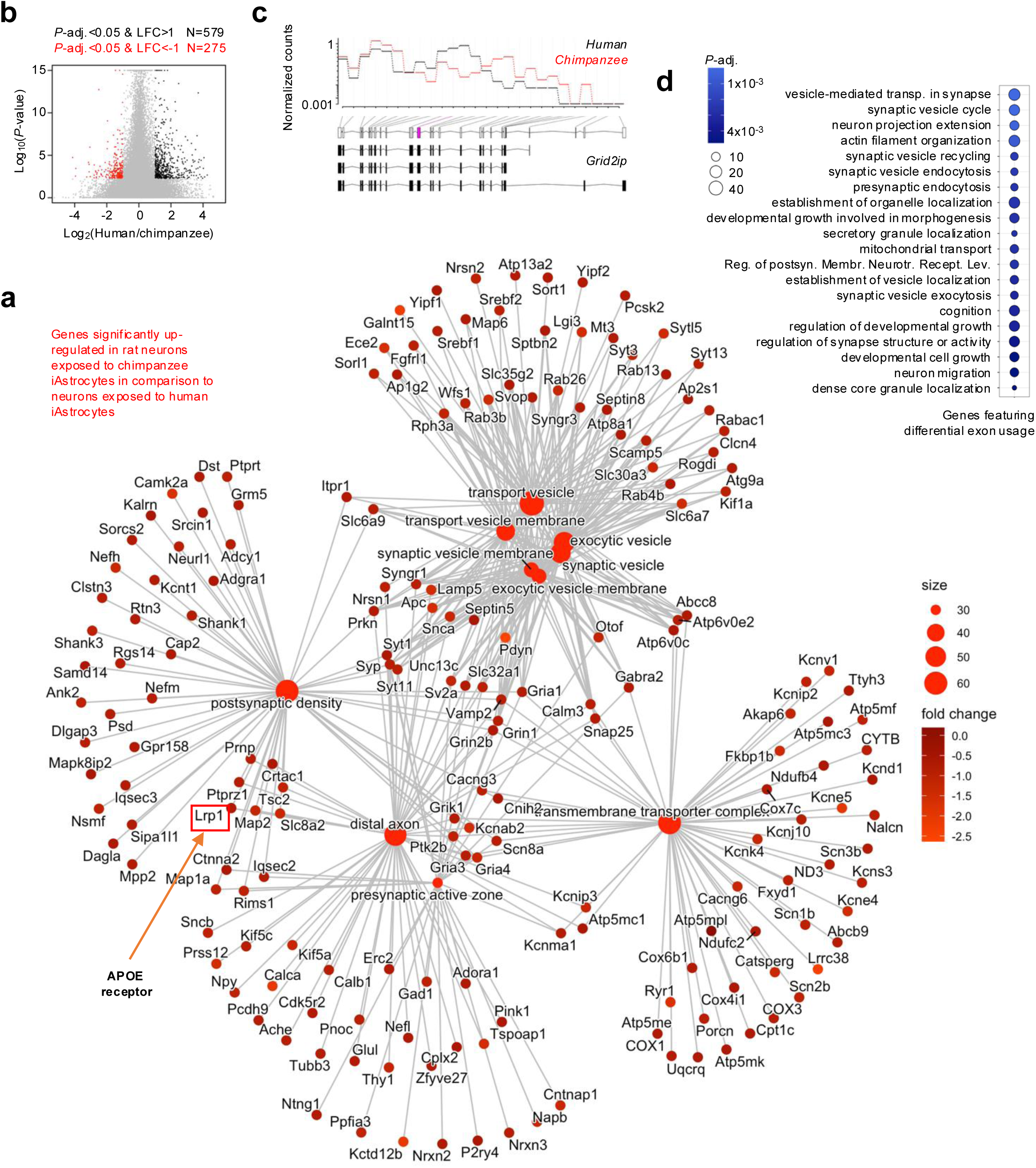
Astrocyte-dependent regulation of neuronal maturation programs at transcriptional and splicing levels. **a** Gene Ontology network representation of GO Cellular Component (GO: CC) categories enriched among genes upregulated in rat neurons co-cultured with chimpanzee iAstrocytes compared with human iAstrocytes. Differential expression was assessed using *DESeq2* (*design = ∼ batch + species*). Genes with LFC < −0.59 and adjusted *P* < 0.1 were used for enrichment analysis. Nodes represent enriched GO categories (large nodes) and individual genes (small nodes). Node size reflects the number of genes contributing to each category, and gene color indicates log_2_ fold change. The network highlights enrichment of synaptic structures, including synaptic vesicles, synaptic vesicle membrane, exocytic vesicles, postsynaptic density, presynaptic active zone, and distal axon, revealing activation of molecular programs associated with vesicle trafficking and synaptic transmission in neurons exposed to chimpanzee iAstrocytes. The gene encoding APOE receptor LRP1 is highlighted within the postsynaptic density module. **b** Volcano plot showing differential exon expression in rat neurons exposed to human versus chimpanzee iAstrocytes inferred using *DEXSeq*-based analysis. Exons significantly upregulated in neurons exposed to chimpanzee astrocytes (*P*-adj.<0.05, LFC < −1) are highlighted in red, whereas exons significantly upregulated in neurons exposed to human astrocytes (*P*-adj.<0.05, LFC>1) are highlighted in black. **c** Representative differential exon usage profile (*DEXSeq* tool) for *Grid2ip*, illustrating coordinated, astrocyte-dependent shifts in exon inclusion across the gene. Black and red traces correspond to neurons exposed to human and chimpanzee iAstrocytes, respectively. **d** Functional enrichment of genes exhibiting differential exon usage, showing overrepresentation of synaptic and vesicle-related processes. Point size reflects the number of genes per category, and color indicates statistical significance.

**Supplementary Figure 7.**
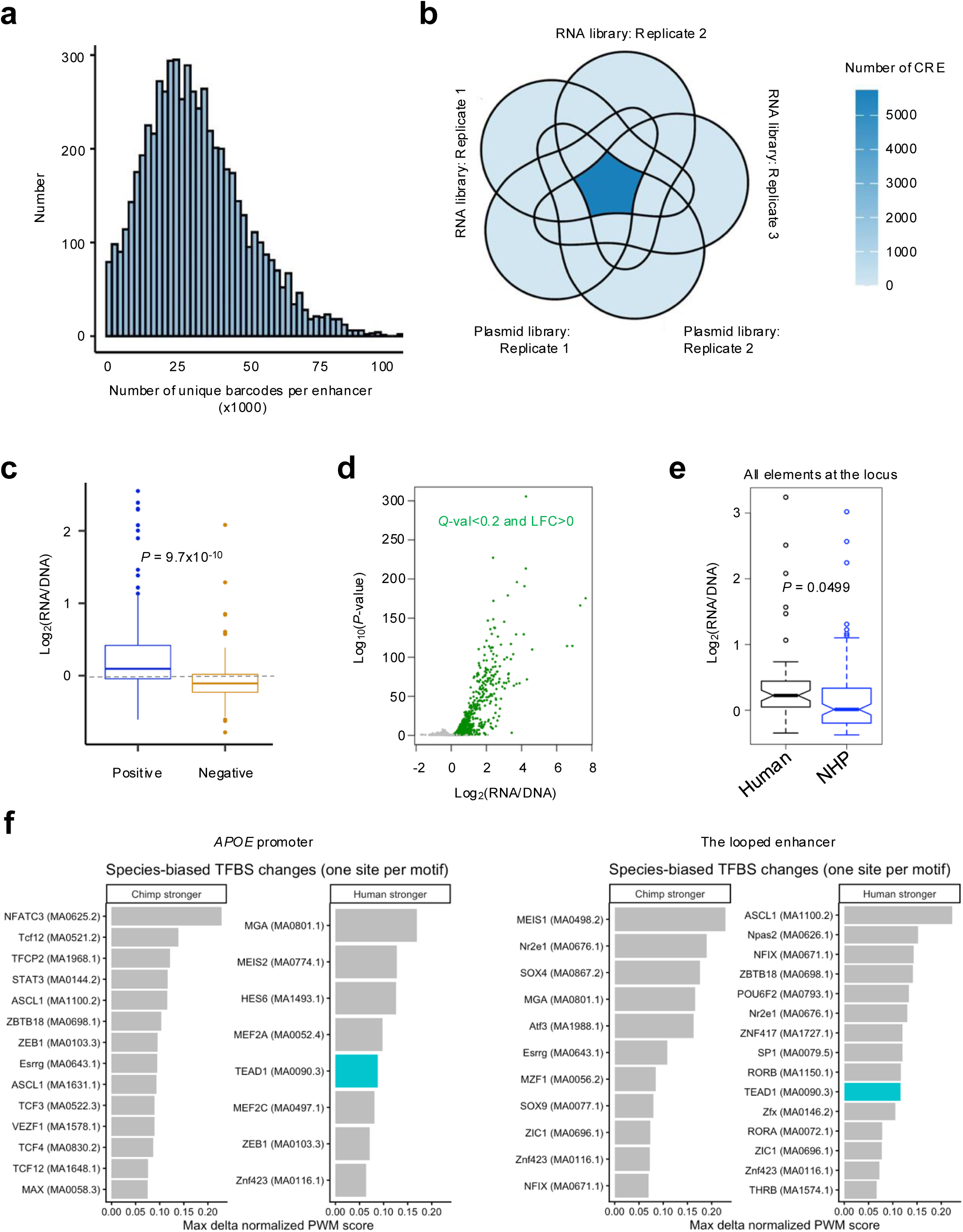
Massively parallel reporter assay (MPRA) identifies regulatory elements at the *APOE* locus. **a** Distribution of barcode representation per candidate cis-regulatory element (CRE) in the MPRA library. The histogram shows the number of unique barcodes associated with each tested element. **b** Overlap of CREs detected across MPRA replicates. The Venn diagram illustrates the number of elements detected in two plasmid library replicates and three RNA library replicates. **c** MPRA activity of known enhancer elements (Positive) and a set of negative control elements (sequences that did not overlap ATAC-seq peaks). Elements regarded as positive control featured a significantly higher MPRA activity than elements from the Negative control set (*P* – two-sided t-test). **d** Identification of active CREs. The scatter plot shows the relationship between CRE activity (log_2_(RNA/DNA)) and statistical significance of difference in barcode abundance within the RNA and DNA sequencing libraries (−log_10_(*P*-value), *DESeq2* test). Green points represent elements that pass the significance thresholds (q-value < 0.2 and log_2_(RNA/DNA) > 0) and are therefore termed active CREs. **e** Comparison of regulatory activity between human and non-human primate (NHP) sequences across all tested elements at the *APOE* locus (+/- 500 kb around APOE promoter). Boxplots display the distribution of enhancer activity log_2_(RNA/DNA). NHP sequences showed significantly higher activity compared with human sequences (*P* = 0.0499, two-sided t-test). **f** Species-biased changes in predicted TF motif strength in the *APOE* promoter (left) and enhancer (right). A 1 kb orthologous enhancer region was analyzed in human and chimpanzee using reciprocal motif scanning with JASPAR vertebrate PWMs. In the human-centered analysis, candidate TF binding sites were first identified in the human sequence, and the orthologous chimpanzee sequence was then extracted from a pairwise human-chimpanzee alignment. In the chimpanzee-centered analysis, candidate sites were first identified in the chimpanzee sequence, and the orthologous human sequence was extracted analogously. For each matched site, raw PWM scores were recalculated on the exact human and chimpanzee sequences and converted to motif-specific normalized scores using the PWM minimum-to-maximum score range. Motif score differences were then calculated as human-chimpanzee in the human-centered analysis and chimpanzee-human in the chimpanzee-centered analysis. Only non-indel-disrupted sites with at least one nucleotide difference between species that featured a normalized score greater than 0.84 in at least one species, an absolute normalized score difference greater than 0.05, and TFs expressed in astrocytes were retained. In the summary plot, the strongest site per motif was retained for the human-stronger and chimpanzee-stronger sets. TEAD-family motifs are highlighted in turquoise; all other motifs are shown in gray.

**Supplementary Figure 8.**
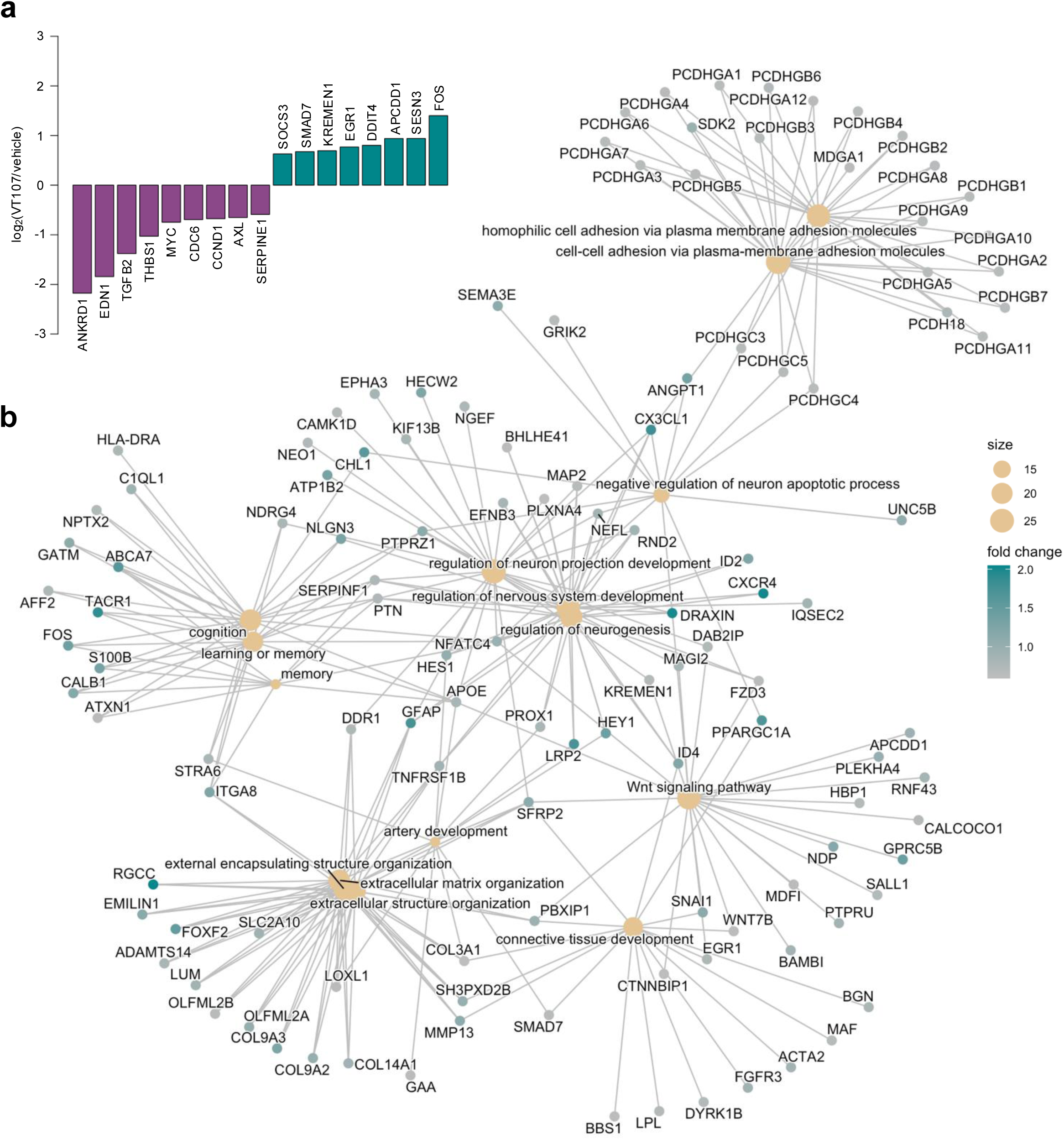
TEAD inhibition remodels astrocyte transcriptional programs. **a** Expression changes of representative YAP/TAZ-TEAD target genes following pharmacological inhibition of TEAD auto-palmitoylation using VT-107 in human iAstrocytes. The bar plot shows log_2_ fold change relative to vehicle-treated controls. Canonical TEAD-responsive genes involved in growth factor signaling and proliferative programs (e.g., ANKRD1, EDN1, TGFB1, CCN1, CCN2) are downregulated, whereas stress-response and signaling feedback regulators (e.g., SOCS3, SMAD7, KREMEN1, DDIT4, FOS) are induced. **b** Gene ontology network representation of biological processes enriched among genes upregulated following TEAD inhibition in human iAstrocytes. Nodes represent enriched GO biological process terms (large nodes) and associated genes (small nodes). Node size reflects the number of genes contributing to each category, and gene color indicates fold change.

